# Identification of Human Immune Cell Subtypes Most Vulnerable to IL-1β-induced Inflammatory Signaling Using Mass Cytometry

**DOI:** 10.1101/2020.04.19.047274

**Authors:** Hema Kothari, Corey M. Williams, Chantel McSkimming, Mythili Vigneshwar, Eli R. Zunder, Coleen A. McNamara

## Abstract

IL-1β has emerged as a key mediator of the cytokine storm linked to high morbidity and mortality from COVID-19 and blockade of the IL-1 receptor (IL-1R) with Anakinra has entered clinical trials in COVID-19 subjects. Yet, knowledge of the specific immune cell subsets targeted by IL-1β and IL-1β-induced signaling pathways in humans is limited. Utilizing mass cytometry (CyTOF) of human peripheral blood mononuclear cells, we identified effector memory CD4 T cells and CD4^−^CD8^low/-^CD161^+^ T cells as the circulating immune subtypes with the greatest expression of p-NF-κB in response to IL-1β stimulation. Notably, CCR6 distinctly identified T cells most responsive to IL-1β. Other subsets including CD11c myeloid dendritic cells (mDCs), classical monocytes (CM), two subsets of natural killer cells (CD16^−^CD56^bright^CD161^−^ and CD16^−^CD56^dim^CD161^+^) and a population of lineage^−^(Lin-) cells expressing CD161 and CD25 also showed IL-1β-induced expression of p-NF-kB. The IL-1R antagonist, Anakinra significantly inhibited IL-1β-induced p-NF-kB in the CCR6^+^ T cells and CD11c mDCs with a trending inhibition in CD14 monocytes and Lin^−^CD161^+^CD25^+^ cells. IL-1β also induced a rapid but much less robust increase in p-p38 expression as compared to p-NF-kB in the majority of these same immune cell subsets. Prolonged IL-1β stimulation greatly increased p-STAT3 and to a much lesser extent p-STAT1 and p-STAT5 in T cell subsets, monocytes, DCs and the Lin^−^CD161^+^CD25^+^ cells suggesting IL-1β-induced production of downstream STAT-activating cytokines, consistent with its role in cytokine storm. Interindividual heterogeneity and inhibition of this activation by Anakinra raises the intriguing possibility that assays to measure IL-1β-induced p-NF-kB in CCR6^+^ T cell subtypes could identify those at higher risk of cytokine storm and those most likely to benefit from Anakinra therapy.

## INTRODUCTION

Cytokine storm is suspected of producing the overzealous immune response driving the severe cardiopulmonary complications that are fueling the high morbidity and mortality rates in certain COVID-19-infected individuals ^1,2^. Cytokine storm or macrophage activation syndrome or secondary haemophagocytic lyphohistocytosis has previously been described as a serious complication in other clinical contexts including individuals with rheumatologic disorders such as systemic onset juvenile inflammatory arthritis also known as Still’s disease ^3^, sepsis ^4^, and CAR-T cell therapy ^5^. IL-1β is one of the key inflammatory cytokines implicated in the cytokine storm syndrome ^4,6,7^. Inhibition of IL-1β-induced cellular signaling with the IL-1 receptor (IL-1R) antagonist, Anakinra, has shown promise in treating cytokine storm and is proposed as a potential therapy for subjects with this severe complication of COVID-19 ^8–10,11^. Yet, little is known about the immune cell subtypes most vulnerable to IL-1β-induced inflammatory signals in humans. New technologies, such as mass cytometry (CyTOF), have emerged as powerful analytic tools to identify unique cell phenotypes associated with specific inflammatory stimuli and their inhibitors ^12–14^. CyTOF uses antibodies conjugated to isotopes of rare-earth metals not found in human cells, so it eliminates background signal and minimizes signal overlap, allowing for the identification of a greater number of proteins that more precisely define cellular identify and activation states. These novel, customized approaches hold promise for the discovery of the cellular mechanisms mediating IL-1β-induced pathologies and the identification of sensitive and specific biomarkers that could be used to assist in early identification of cytokine storm and those who may response to specific therapies such as IL-1β or IL-1R inhibitors.

NF-κB, p38 MAPK, ERK and JNK activation have been implicated in IL-1R1-mediated immune cell activation ^15–23^. Notably, the majority of these are murine studies. Based on these findings, we developed a customized 35 antibody IL-1β-response CyTOF panel to generate the first comprehensive atlas of IL-1β-induced signaling in peripheral blood mononuclear cells (PBMCs) in humans. Results demonstrated that IL-1β rapidly induces p-NF-kB and p-p38 in distinct effector memory (EM) T subsets, monocytes, dendritic cells (DC), natural killer (NK) cells and lineage-(Lin^−^)CD161^+^CD25^+^ cells. IL-1β-induced increase in p-p38 was much less robust than p-NF-kB. In contrast to the rapid increase of p-NF-kB with only 15 minutes of IL-1β stimulation, prolonged stimulation for one hour greatly increased p-STAT3 and to a lesser extent p-STAT1 and p-STAT5 in subsets of T cells, monocytes, DCs and Lin^−^CD161^+^CD25^+^ cells. Notably, we identified CCR6 as a marker to distinctly identify T cells most responsive to IL-1β. The IL-1R antagonist Anakinra significantly inhibited IL-1β-induced p-NF-kB in the CCR6^+^ T cells and CD11c myeloid DCs (mDC) with only a trending inhibition in CD14 monocytes and Lin^−^CD161^+^CD25^+^ cells. This knowledge of the cell types, unique markers enriched on responsive cells and temporal expression of inflammatory signaling molecules in response to IL-1β stimulation in humans provides unique information that may be used to discover new approaches to limit IL-1β-induced cytokine storm and other pathologies. Additionally, the heterogeneity in subject response to IL-1β suggests that findings could be utilized to identify those most at risk for cytokine storm and those most likely to benefit from IL-1β inhibition.

## MATERIALS AND METHODS

### Healthy Donors

Peripheral blood from healthy volunteers was obtained after written informed consent under the guidelines of the Institutional Review Board (IRB protocol # 16017).

### PBMC Isolation

Blood from healthy donors was drawn into BD K2 EDTA vacutainer tubes and processed at room temperature (RT) within one hour of collection. Whole blood in vacutainers were centrifuged at 400 x g for 10 minutes at RT to remove platelet rich plasma. PBMCs were separated by Ficoll-Paque density-gradient centrifugation (Ficoll-Paque PLUS (GE Healthcare Biosciences AB) and SepMate-50 (Stemcell Technologies Inc) following the manufacturer’s protocol. Trypan blue staining was performed to get live cell counts. PBMCs were cryopreserved in freezing solution (90% FBS/10% DMSO). PBMC vials were stored in Mr. Frosty (Thermo Fisher) for 48 hours at −80°C and were then stored in liquid nitrogen until used.

### CyTOF Panel Optimization

All metal-conjugated antibodies were purchased from Fluidigm and purified unlabeled antibodies from the vendors as shown in **Supplementary Table 1**. Unlabeled antibodies were conjugated in-house using the MaxPAR antibody labeling kit (Fluidigm) according to the manufacturer’s protocol. After determining the percent yield by measurement of absorbance at 280 nm, metal-labeled antibodies were diluted in Candor PBS Antibody Stabilization solution (Candor Bioscience GmbH) for long-term storage at 4°C.

Cryopreserved PBMCs were used for CyTOF panel optimization. Antibody titrations were perfomed to determine optimal antibody concentrations. For titration of phospho signaling antibodies, PBMCs were stimulated with PMA/Ionomycin (50 nM/1 µg/ml) for 15 minutes. To capture IL-1β-induced transient signaling, PBMCs needed to be fixed immediately with paraformaldehyde (PFA, 1.6% final concentration) post stimulation. As such, impact of PFA on integrity of surface marker epitopes was evaluated. PFA fixation led to significant loss of CXCR4, CXCR5, CCR6, CCR7 and CD127 staining while increased non-specific binding of CXCR3 (**Supplementary Fig. 1**). As such, an alternative approach was utilized to label cells with these PFA-labile antibodes. A small volume of a cocktail of these PFA-labile antibodies was added simultaneously during IL-1β stimulation of PBMCs prior to PFA fixation. Antibody addition during PBMC stimulation did not interfere with IL-1β-mediated signaling (**Supplementary Fig. 2**), while allowed specific staining. Preliminary analysis did not show expression of p-JNK and ribosomal protein p-S6, an AKT effector upon IL-1β stimulation so these markers were excluded from the panel.

### PBMC Stimulation, Barcoding and CyTOF Staining

Cryopreserved PBMCs were thawed and washed twice with warm complete media (RPMI supplemented with 5% FBS, 1 mM sodium pyruvate and Pen-Strep). All PBMC samples had viability in the range of 85-98%. 1-2 × 10^6^ cells per stimulation were used. Prior to IL-1β (Peprotech, Rocky Hill, NJ) stimulation, PBMCs were rested in 1.8 ml low retention eppendorf tubes (Fisherbrand) for 1 hour at 37°C in 5% CO_2_. At the end of the stimulation period, PFA (1.6% final concentration) methanol free, Ultra Pure EM grade, Polysciences, Inc) was immediately added to the PBMC suspension, mixed by vortexing, and cells were incubated at RT for 10 minutes. PFA was then diluted at least to 0.8% or lower with ice-cold Maxpar Cell Staining Buffer (CSB) (Fluidigm), and fixed cells were centrifuged at 800 × g for 10 minutes at RT. PBMC samples were barcoded using the palladium-based 20-Plex Pd Barcoding Kit (Fluidigm) according to the manufacturer’s protocol. As PBMCs were fixed with PFA, which is similar to the Maxpar Fix I Buffer, this step was excluded and cells were washed twice with 1 ml of 1x Maxpar Barcode Perm Buffer. Barcoded cells were then combined into a single tube prior to Fc receptor blocking (BD Biosciences) and staining with a cocktail of metal-conjugated antibodies against cell-surface markers for 30 minutes at RT with frequent gentle shaking. After washing once with the CSB, cells were resuspended in residual CSB and chilled on ice for 5 minutes. For cell permeabilization, 0.5 ml of 100% ice-cold methanol was added to the cells, and samples were vortexed and incubated on ice for 10 minutes. Eppendorf tubes were filled up with the CSB and samples were centrifuged at 800 × g for 10 minutes at RT. An additional wash with the CSB was performed to remove residual methanol prior to staining with antibodies against intracellular phospho markers for 45 minutes at RT. After washing, cell were fixed with 2% PFA for 10 minutes at RT, washed again and cell pellet stored overnight at 4°C. Next day, stained cells were incubated with iridium DNA intercalator (Fluidigm) in Maxpar Fix and Perm Buffer (Fluidigm) for 20 minutes at RT. Prior to acquisition, cells were washed once with CSB followed by two washes with the Maxpar Cell Acquisition Solution (Fluidigm). Cells were filtered through a 40 μm membrane just before being acquired on a Helios mass cytometer (Fluidigm). EQ Four Element Calibration Beads from Fluidigm were added as per the manufacturer’s directions prior to running.

### CyTOF data pre-processing and subsequent analysis

Data obtained from the Helios instrument were in the form of. fcs files format. Samples were normalized using the Nolan lab MATLAB normalizer (http://github.com/nolanlab/bead-normalization/releases) and debarcoded using Zunder’s lab debarcoder ^24^ (https://github.com/zunderlab/single-cell-debarcoder). Normalized and debarcoded. fcs files were then analyzed in Flowjo version 10.3 for Mac (Flowjo LLC, Ashland, OR). Arcsinh transformation was performed on all measurements ^13^. Clean-up gating was done on barcode stringency parameters and iridium DNA intercalator to remove non-cell debris and cellular aggregates (**Supplementary Fig. 3a,b**). The. fcs files were uploaded to Cytobank for Spanning-tree Progression Analysis of Density-normalized Events (SPADE). For SPADE tree construction, clean CD45^+^ cells and cell surface markers as shown in **Supplementary Table 1** were used for clustering using the default settings in Cytobank.

Cell types were identified by first manually gating total CD3^+^ (for T cells), CD3^−^CD19^−^CD56^−^HLA-DR^+^ cells (for monocytes and DCs) and CD3^−^CD19^−^HLA-DR^−^cells (for NK cells) (**Supplementary Fig. 3c-e**), and then passing through an R pipeline similar to Nowicka et al ^25^. Dimensionality reduction by UMAP ^26,27^ was performed in R using the “UMAP” package. All parameters were used at default values except for min_dist, which was changed to 0.01. Leiden clustering ^28^ was performed on the UMAP graph using Modularity Vertex Partition in the Python Leidenalg package. Cell surface markers used for UMAP and Leiden clustering for each individual immune cell subsets (T cells, monocytes/DCs and NK cells) are described in the results. Cells were gated in Flowjo to quantitate frequency of cells expressing phospho markers within immune cell clusters (**Supplementary Fig. 3f-j**).

### Statistics

GraphPad Prism Version 8.2.0 (GraphPad Software, Inc) was used for data analysis, graphing and statistics. All data are expressed as mean ± SEM and analyzed with the 2-tailed parametric paired or unpaired student *t* test. A *p*<0.05 was considered statistically significant.

## RESULTS

### IL-1β induced early increases in p-NF-kB and p-p38 in distinct immune cell subtypes in humans

A 35 antibody panel was developed based on a comprehensive review of the literature on IL-1β-induced effects on immune cells ^15–23^ (Fig. 1a and **Supplementary Table 1**) and optimized (**Supplementary Figs. 1,2**). The panel included antibodies to major immune cell types, IL-1Rs and signaling proteins including p-NF-kB, p-p38, p-ERK, p-STAT1, p-STAT3 and p-STAT5. PBMCs from human donors were stimulated with vehicle or IL-1β for 5, 15, 30, 60, 120 and 240 minutes, fixed, barcoded and analyzed by CyTOF with subsequent computational data analysis (Fig. 1b). PBMCs stimulated with varying doses of IL-1β (10, 25 and 100 ng/ml) revealed equivalent responses in terms of expression of p-NF-kB and p-p38 (**Supplementary Fig. 4**), so 10 ng/ml dose was used for all subsequent experiments. To map IL-1β-induced signaling responses across major immune cell types, Spanning-tree Progression Analysis of Density-normalized Events (SPADE) analysis was applied on all vehicle and IL-1β-stimulated samples. SPADE analysis identified 9 major immune cell populations including, CD4^+^, CD8^+^, and CD4^−^CD8^low^ T cells, CD19^+^ B cells, HLA-DR^+^CD14^+^ and HLA-DR^+^CD16^+^ monocytes, HLA-DR^+^CD11c^+^ (mDCs) and HLA-DR^+^CD123^+^ plasmacytoid DCs (pDCs), and HLA-DR^−^CD56^+^CD16^+^ NK cells (Fig. 1c,d). CD4 T cells are the most abundant cell type within PBMCs (Fig. 1d). The representative SPADE tree shows IL-1β-induced expression of p-NF-kB, p-p38 and p-ERK across different immune cell types with 15 minutes of stimulation (Fig. 1e). This time point was chosen for this representative figure and deeper immune subtype analysis in figures 2–6 as time course analysis demonstrated IL-1β-induced direct signaling activation most robust at this time point (**Supplementary Fig. 5a,b**). While T cells, monocytes and DCs in the vehicle-treated samples showed basal levels of p-NF-kB, p-p38 and p-ERK, IL-1β stimulation led to a marked upregulation of p-NF-kB expression prominently in the CD4 and CD4^−^CD8^low^ T cell subsets (Fig. 1e). Additionally, a modest level of IL-1β-induced p-NF-kB expression was observed in the CD11c mDCs and NK cells. IL-1β induced p-p38 expression primarily in CD14 monocytes and to a lower extent in CD11c mDCs and CD4 T cells. IL-1β did not induce p-ERK expression in any immune cell type, and B cells were relatively unresponsive to IL-1β stimulation (Fig. 1e). As expected, short-time stimulation of PBMCs with IL-1β did not induce STAT activation (Fig. 6a and **Supplementary Fig. 6**).

**Fig 1.**
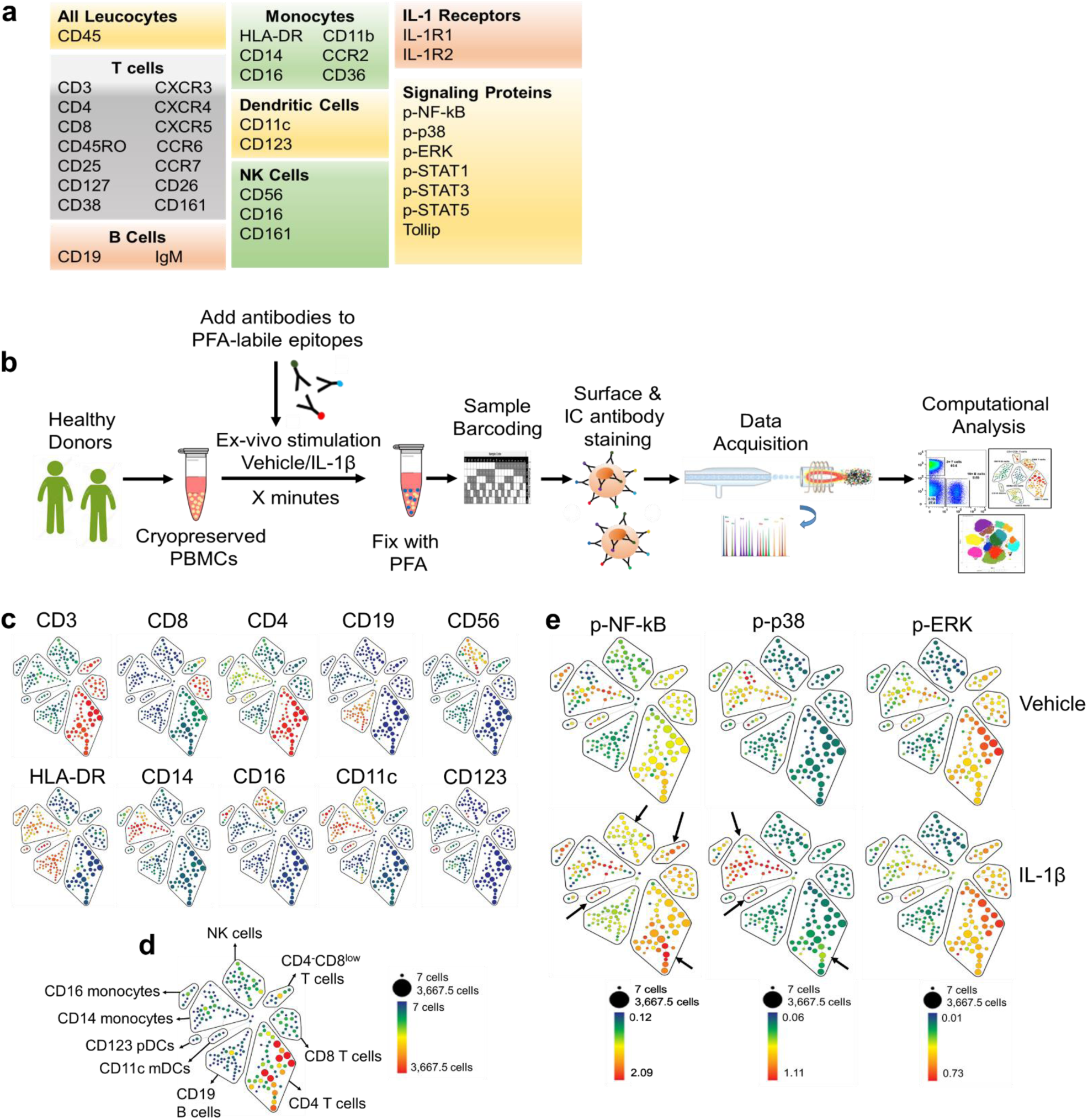
IL-1β induces rapid expression of p-NF-kB and p-p38 in distinct immune cell subtypes. **a**, CyTOF antibody panel. Note: markers including chemokine receptors, CD161, CD11b, CD11c and others are expressed on more than one cell type. **b**, Experimental design. **c**, SPADE analysis was performed using cell surface markers to identify 200 cell clusters using 10% down-sampling. SPADE tree showing median expression (blue to red depicting low to high) of the indicated lineage markers. The size of each node is proportional to the number of cells in that population. **d**, 9 immune cell types were identified: Total T cells were identified as CD3^+^, and subsets as CD4^+^, CD8^+^ and CD4^−^CD8^low^, B cells as CD19^+^, NK cells as CD56^+^CD16^+^, monocyte subsets as HLA-DR^+^CD14^+^ and HLA-DR^+^CD16^+^ and DCs as HLA-DR^+^CD11c^+^ and HLA-DR^+^CD123^+^. Scale: Color blue to red within each cluster corresponds to low to high cell count. **e**, Representative SPADE tree from a donor showing IL-1β-induced expression of p-NF-kB, p-p38 and p-ERK. Note: the scale min and max depicting median expression differ for each phospho marker.

**Fig 2.**
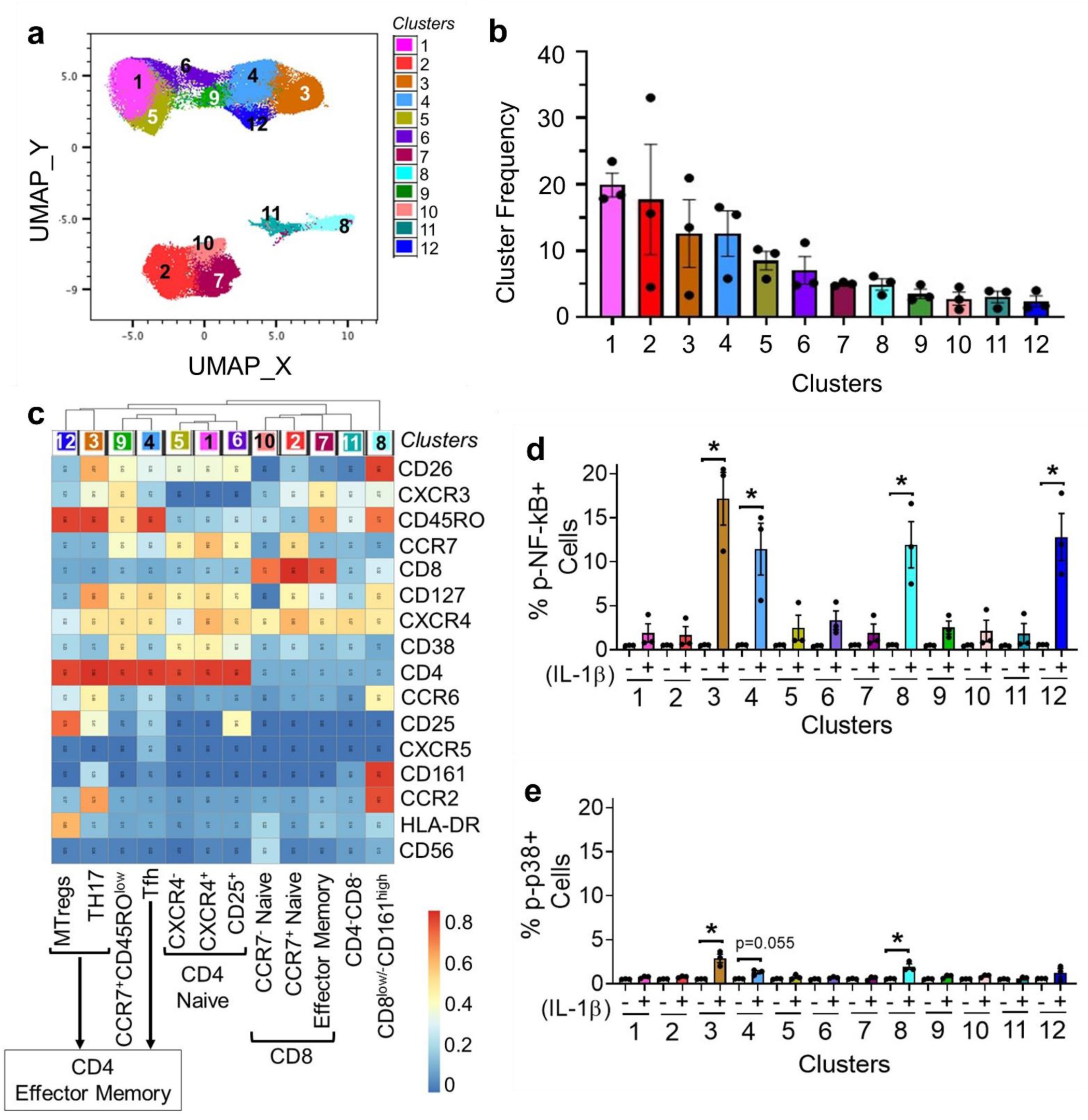
Effector-memory T cell subsets are the predominant subsets that express IL-1β induced p-NF-kB and p-p38. UMAP followed by Leiden clustering was performed on CD3^+^ T cells using both vehicle and IL-1β-stimulated cells and cell surface markers as shown in **c**. **a**, 12 T cell clusters were identified as shown superimposed on the UMAP. **b**, Frequency of T cell clusters and inter-individual variations between donors. **c**, Heatmap showing median expression of surface markers across all 12 T cell clusters. The color in the heatmap represents the median of the arcsinh for each subset with 0-1 transformed marker expression. The dendrogram represents the hierarchical similarity for subsets. Frequency of **d**, p-NF-kB^+^ cells and **e**, p-p38^+^ cells within each cluster in the vehicle-treated and IL-1β-stimulated samples. Significant differences represent the comparisons between vehicle and IL-1β-stimulated samples. Data shown is mean ± SEM. Paired t-test. N=3; * p-value ≤ 0.05.

### Effector memory T cells show the strongest increase in expression of p-NF-kB and p-p38 in response to IL-1β stimulation

To identify specific T cell subsets that show IL-1β-induced increase in p-NF-kB and p-p38, we performed Uniform Manifold Approximation and Projection (UMAP), followed by Leiden clustering on CD3^+^ T cells (**Supplementary Fig. 3d**). Unsupervised clustering of CD3^+^ T cells resulted in 12 clusters (Fig. 2a). The frequency of each cluster is shown in Fig. 2b. Clusters were annotated based on the expression of lineage markers as shown in the heatmap in Fig. 2c. A marked increase in the frequency of p-NF-kB^+^ cells was observed in TH17 cells (cluster 3; CD4^+^CD45RO^+^CCR6^+^CD161^+^CD26^+^), T follicular helper cells (Tfh) (cluster 4; CD4^+^CD45RO^+^CXCR5^+^), CD8^low/-^CD161^+^ T cells (cluster 8) and memory T regulatory cells (Mtregs) (cluster 12; CD4^+^CD45RO^+^CD25^+^CD127^low/-^) (Fig. 2d). Cluster 8 showed a bimodal expression of CD8 and biaxial gating demonstrated presence of two distinct cell populations within cluster 8: CD8^−^(57.6 ± 2.00%; SEM) and CD8^+^ (40.5 ± 2.16%; SEM) that were both CD4^−^CD45RO^+^, expressed comparable levels of CD161, and showed IL-1β-induced expression of p-NF-kB (**Supplementary Fig. 7a-c and i**). Moreover, similar to TH17 cells (cluster 3), both CD8^−^ and CD8^+^ subpopulations within cluster 8 expressed CD26 (**Supplementary Fig. 7d**), a marker that is associated with cells capable of producing IL-17 ^29^. Notably, all four IL-1β responsive T cell clusters were negative for CCR7 (Fig. 2c), indicating that IL-1β specifically targets the EM T cells in humans. Similar to p-NF-kB, IL-1β-induced expression of p-p38 was observed in TH17 cells (cluster 3), Tfh cells (cluster 4) and CD8^low/-^CD161^high^ cells (cluster 8) (Fig. 2e) however, there was a much lower frequency of cells expressing p-p38 as compared to p-NF-kB (1.5-3.5% vs 5-20%). Notably, there was clear variation (5-20%) in IL-1β-induced expression of p-NF-kB in these T cell subtypes between individuals suggesting that this assay may have the potential to identify individual differences in susceptibility to IL-1β-induced downstream effects.

### CCR6 expression distinctly identifies T cells most vulnerable to IL-1β

As IL-1β signals through binding to its surface receptor IL-1R1, we reasoned that the memory T cells that show induced expression of p-NF-kB or p-p38 upon IL-1β stimulation express IL-1R1 while the remaining memory T cells do not. However, mean expression of IL-1R1 was significantly lower in the p-NF-kB^+^ memory T cells in comparison to the p-NF-kB^−^memory T cells (Fig. 3a). IL-1β signaling can induce receptor internalization ^30^ yet, there was no reduction in the mean expression of surface IL-1R1 after 15 minutes of IL-1β stimulation in both the p-NF-kB^−^ and p-NF-kB^+^ memory T cell subsets (Fig. 3a). Cell surface IL-1R2 binds to IL-1β but acts as a decoy receptor ^31^. As such, the ratio of the levels of IL-1R1 and IL-1R2 on the cell surface may determine the signaling outcome induced by IL-1β. Therefore, we also analyzed expression of IL-1R2 on the p-NF-kB^+^ and p-NF-kB^−^ memory T cells. IL-1R2 expression could not be detected on T cells or any other immune cell type in both vehicle-treated and IL-1β-stimulated PBMCs. Notably, the same IL-1R2 antibody clone measured IL-1R2 expression on CD4 memory T cells activated with anti-CD3, anti-CD28 and IL-2 for 48 hours. Unstimulated cells in culture also showed moderate levels of IL-1R2 expression, yet it was undetectable on unstimulated freshly purified PBMCs (**Supplementary Fig. 8**).

**Fig 3.**
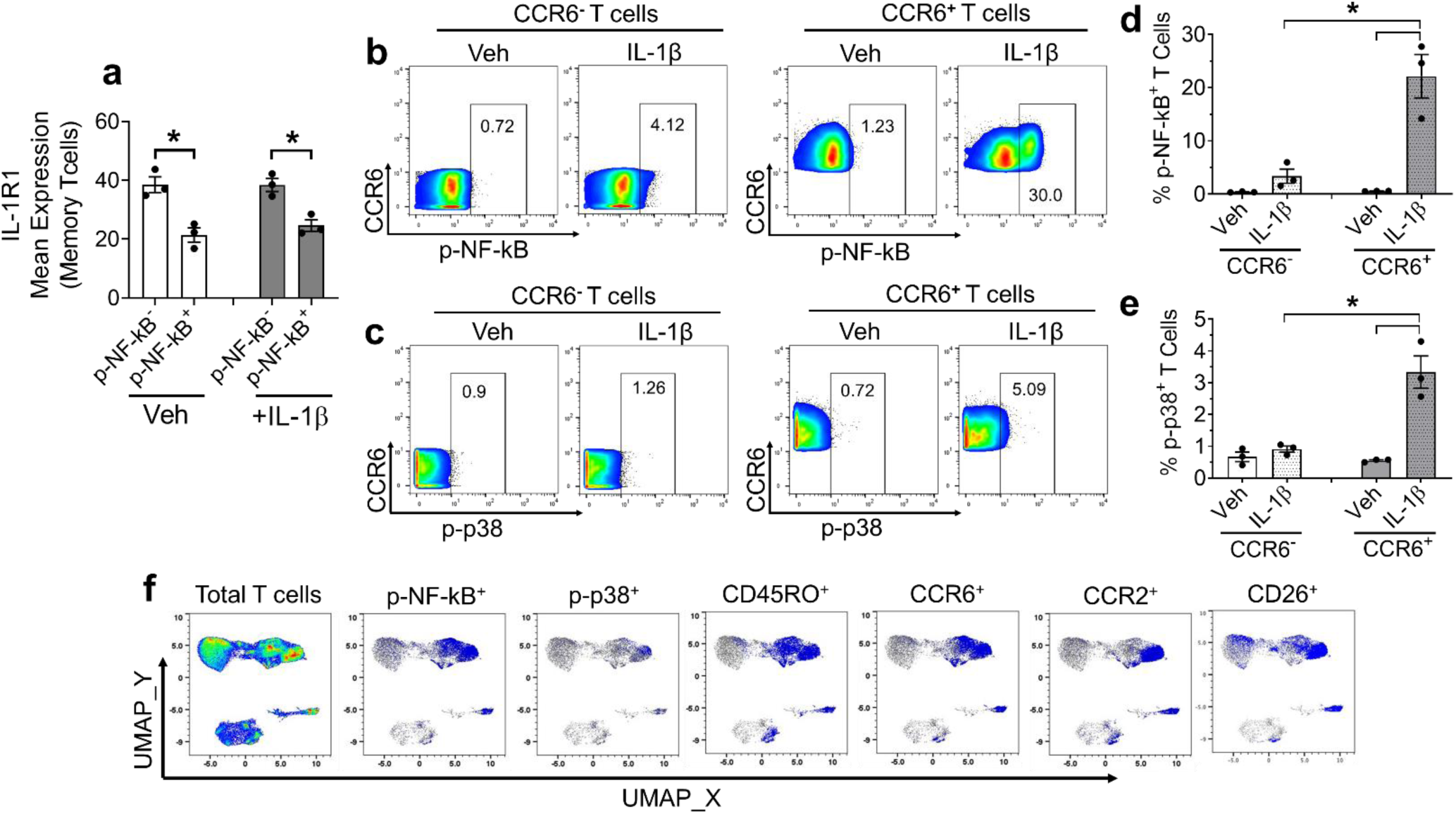
CCR6 expression marks T cells that are most vulnerable to IL-1β. CD3^+^ memory T cells were separated into p-NF-kB- and p-NF-kB^+^ subpopulations in the vehicle-treated and IL-1β-stimulated samples. **a**, Mean expression of IL-1R1 within the p-NF-kB- and p-NF-kB^+^ memory T cells. **b and c**, Representative dot blots showing frequency of p-NF-kB^+^ and p-p38^+^ cells within CCR6^−^ and CCR6^+^ T cell subsets in the vehicle-treated and IL-1β-stimulated samples. **d and e**, Quantified frequency of p-NF-kB^+^ and p-p38^+^ cells. **f**, UMAP overlays showing co-localization of p-NF-kB^+^ and p-p38^+^ cells specifically within the CD45RO^+^ memory T cell subset that expresses CCR6. Moreover, CCR2^+^ and CD26^+^ T cells superimposed on the UMAP show that a majority of the CCR6^+^ memory T cells that show IL-1β-induced signaling co-express CCR2 and CD26. Data shown is mean ± SEM (Unpaired t-test. N=3; * p-value ≤ 0.05).

Notably, as shown in Fig. 2c, CCR6 was the only marker that was uniquely expressed in the four T cell clusters (3, 4, 8 and 12) that showed IL-1β-induced expression of p-NF-kB. Yet, unlike lineage markers such as CD4, CD8 and CD45RO, CCR6 median expression was moderate, raising the interesting possibility that there may be a range of CCR6 expression in these T cell subtypes, and that CCR6 may mark those most prone to activation in response to IL-1β. To test this, we gated total CD3 T cells into CCR6^−^ and CCR6^+^ populations and analyzed expression of p-NF-kB and p-p38 in these subsets. Indeed, remarkably higher frequencies of p-NF-kB^+^ and p-p38^+^ T cells were present within the T cells that expressed CCR6 as compared to those that do not (Fig. 3b-e). Moreover, the majority (61.5 ± 2.75%) of the CCR6^+^ memory T cells that showed IL-1β-induced expression of p-NF-kB or p-p38 co-expressed CCR2 and CD26 (Fig. 3f).

### Classical monocyte subsets and CD11c myeloid dendritic cells show IL-1β-induced expression of p-NF-kB and p-p38

To identify monocyte subsets and DCs that respond to IL-1β-induced signaling, we used CD3^−^CD19^−^CD56^−^HLA-DR^+^ cells (**Supplementary Fig. 3e**) for UMAP and Leiden clustering. Twelve clusters were identified (Fig. 4a-c). Based on expression of lineage markers, seven clusters (1-6 and 9) belonged to the CM subtypes. Monocyte clustering with markers as shown in Fig. 4c did not separate non-classical (NCM) and intermediate monocytes (IMM). Both NCM and IMM and some CD14^−^CD16^−^ cells were present within cluster 7 as confirmed by biaxial gating (**Supplementary Fig. 9**). Cluster 11 shared a similar phenotype to cluster 7 but expressed high levels of CD123 and was present in only one of the donors (Fig. 4b). Cluster 8 was CD11c^+^CD123^−^mDCs while cluster 10 was CD11c^−^CD123^+^ pDCs. Notably, cluster 12 was found to be Lin- and only expressed CD38 and a low level of CXCR4. IL-1β induced the most robust expression of p-NF-kB (~13-fold higher as compared to vehicle control) in the CM cluster 1 (Fig. 4d). In addition, a modest 2.5 to 5-fold induction in the frequency of p-NF-kB^+^ cells was observed in the other CM clusters including 2, 6 and 9. Interestingly, of all the CM clusters, cluster 1 showed the highest median expression of CCR2 and Tollip, and was distinct from the other CM clusters in terms of lower expression of CD11b and CD45RO (Fig. 4c). Of the two DC clusters, IL-1β-induced an increase in the frequency of p-NF-kB^+^ cells (13 to 30-fold) within the CD11c mDCs (cluster 8) (Fig. 4d). Despite the dramatic fold change, the increase was not statistically significant, likely due to donor heterogeneity. A modest and trending increase in the frequency of p-p38^+^ cells was observed within the same CM clusters (1, 2, 6 and 9) and CD11c mDCs (Fig. 4e). In addition to monocyte subsets and CD11c mDCs, IL-1β also induced expression of p-NF-kB and p-p38 in the HLA-DR^+^Lin-cells (cluster 12).

**Fig 4.**
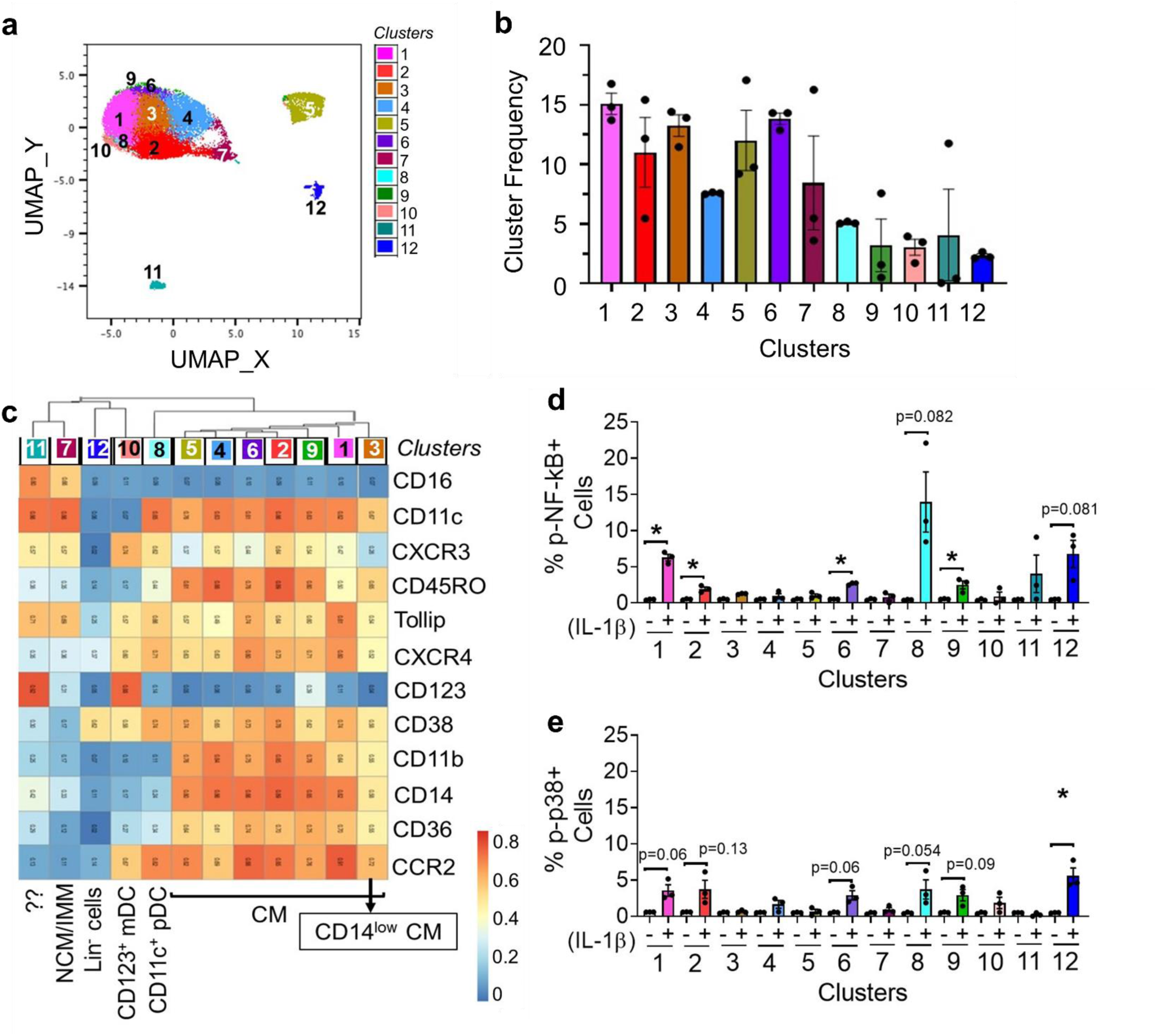
Classical monocytes (CM) and CD11c mDCs show IL-1β-induced expression of p-NF-kB and p-p38. UMAP followed by Leiden clustering was performed on gated CD3^−^CD19^−^CD56^−^HLA-DR^+^ cells using both vehicle and IL-1β-stimulated cells using cell surface markers as shown in **c**. **a**, 12 cell clusters that included monocyte and DC cells were identified as shown superimposed on the UMAP. **b**, Frequency of 12 subsets and inter-individual variations. **c**, Heatmap showing median expression of surface markers across all 12 cell clusters. Frequency of **d**, p-NF-kB^+^ cells and **e**, p-p38^+^ cells within each cluster in the vehicle-treated and IL-1β-stimulated samples. Significant differences represent the comparisons between vehicle and IL-1β-stimulated samples. Data shown is mean ± SEM. Paired t-test. N=3; * p-value ≤ 0.05.

### CD16^−^CD56^−^CD161^+^ cells expressing CD25 show remarkable expression of p-NF-kB in response to IL-1β

CD3^−^CD19^−^HLA-DR^−^ cells (**Supplementary Fig. 3e**) were used for NK cell clustering to identify IL-1β responsive subsets. These CD3^−^CD19^−^HLA-DR^−^ cells included CD56^bright^, CD56^dim^ as well as CD56^−^ populations. In addition to the prototypical NK cell markers such as CD16, CD56 and CD161, other cell surface markers including CXCR3, CXCR4, CD38, CD25, CD11b and CD11c, which were expressed on NK cells were used for the clustering. Interestingly, a total of 17 distinct clusters were identified (Fig. 5a-c). Analysis of IL-1β-induced signaling demonstrated a 10-15-fold induction in the frequency of p-NF-kB^+^ cells in the CD16^−^CD56^dim^CD161^+^ NK cells (cluster 2) and CD16^−^CD56^bright^CD161^−^ (cluster 3) (Fig. 5d). Notably, another CD161^+^ cluster that expressed CD25 but was negative for both CD16 and CD56 (cluster 17) showed the most striking expression of p-NF-kB (>70-fold vs vehicle control) (Fig. 5d). Moreover, IL-1β-induced expression of p-p38 expression was observed only in this cluster (Fig. 5e). Given that CD161 is also expressed on CD4 and CD8 T cell subsets and cluster 17 expressed CD25, we further confirmed that cluster 17 was not contaminating T cells. As shown in the **Supplementary Fig. 10a**, while cluster 17 cells expressed similar levels of CD25 as the T cells, this cluster was negative for CD3, CD4 and CD8 excluding the possibility that these are T cells.

**Fig 5.**
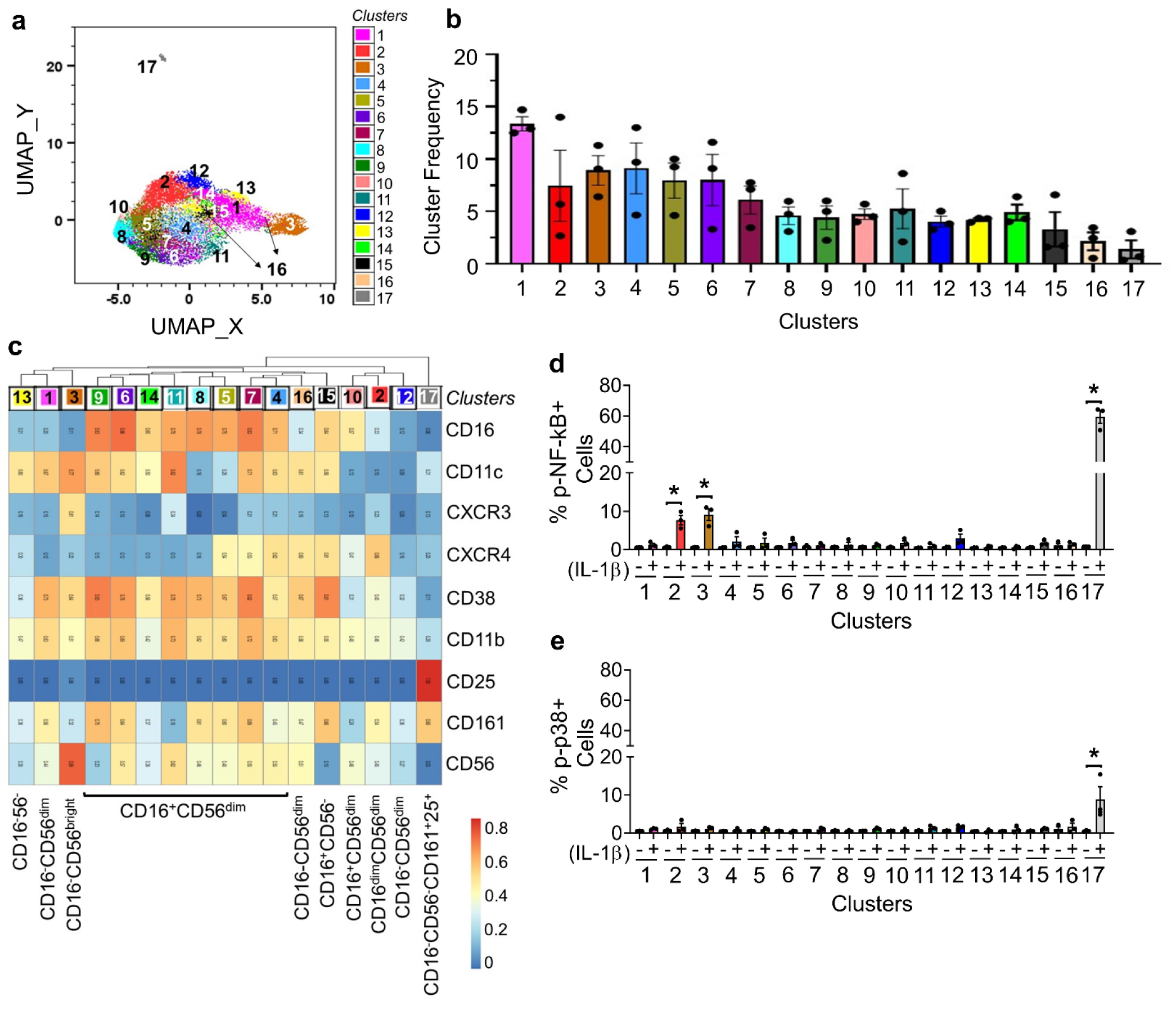
CD16^−^CD56^−^CD161^+^ cells that express CD25 shows remarkable induction of p-NF-kB and p-p38 expression upon IL-1β stimulation. UMAP followed by Leiden clustering was performed on gated CD3^−^CD19^−^HLA-DR^−^ cells using both vehicle and IL-1β-stimulated cells using cell surface markers as shown in **c**. **a**, 17 cell clusters were identified as shown superimposed on the UMAP. **b**, Frequency of clusters and inter-individual variations. **c**, Heatmap showing median intensity of surface markers across all 17 clusters. Frequency of **d**, p-NF-kB^+^ cells and **e** p-p38^+^ cells within each cluster in the vehicle-treated and IL-1β-stimulated samples. Significant differences represent the comparisons between vehicle and IL-1β-stimulated samples. Data shown is mean ± SEM. Paired t-test. N=3; * p-value ≤ 0.05.

### Prolonged IL-1β stimulation induced robust expression of p-STAT3 and to a lesser extent of p-STAT1 and p-STAT5

To identify direct and indirect signaling mediated by IL-1β, time-dependent changes in the IL-1β-induced expression of phospho markers were determined. IL-1β rapidly induced p-NF-kB and p-p38 expression in Th17 cells, Tfh cells, CD8^low/-^CD161^+^ T cells and Mtregs (clusters 3, 4, 8 and 12), CD16^−^CD56^dim^CD161^+^ and CD16^−^CD56^bright^CD161^−^ NK cells (clusters 2 and 3) and Lin^−^CD161^+^CD25^+^ cells (cluster 17) within 15 minutes, which reduced to baseline at later time points (**Supplementary Figs. 5a,b**). Unlike the transient increase in p-NF-kB expression in T and NK clusters, sustained expression of IL-1β-induced p-NF-kB was observed in the CD11c mDCs and CM clusters (**Supplementary Fig. 5a**). Kinetics of p-p38 expression in CD11c mDCs and CM clusters was similar to that of T and NK cells (**Supplementary Fig. 5b**). Prolonged stimulation with IL-1β led to activation of STATs in T cells, monocytes, DCs and Lin_-_CD161^+^CD25^+^ cells (Fig. 6 and Supplementary Fig. 6). Of all the p-STATs analyzed, expression of p-STAT3 was the most robustly induced by IL-1β, particularly in the CD4 naïve and memory T cell clusters and CCR7^+^ CD8 naïve T cells (Fig. 6). Additionally, IL-1β-induced p-STAT3 expression was also observed in CM (clusters 1,2,4,5 and 6), HLA-DR^+^Lin^−^ cells (cluster 12, Fig. 4c), CD11c mDCs, CD123 pDCs, and Lin^−^CD161^+^CD25^+^ cells (cluster 17, Fig. 5c) (**Supplementary Fig. 6a,e**). Notably, p-STAT3 expression in CD123 pDCs was higher as compared to the CD11c mDCs and CM clusters (**Supplementary Fig. 6a**). IL-1β-induced expression of p-STAT1 was seen mostly in the CD4 and CD8 naïve T clusters (**Supplementary Fig. 6g**) and across all monocyte clusters (CM and NC/IMM), CD11c mDCs, CD123 pDCs and Lin^−^CD161^+^CD25^+^ cells (**Supplementary Figs. 6b,d**). As compared to p-STAT3 and p-STAT1, IL-1β induced very low levels of p-STAT5. In contrast to the broad expression of p-STAT5 across monocyte and DC clusters and Lin^−^CD161^+^CD25^+^ cells (**Supplementary Figs. 6c,f**), expression of p-STAT5 in T cells was prominently observed within the Mtregs (cluster 12) and CD4 naïve T cells expressing CD25 (cluster 6) (**Supplementary Fig. 6h**).

**Fig 6.**
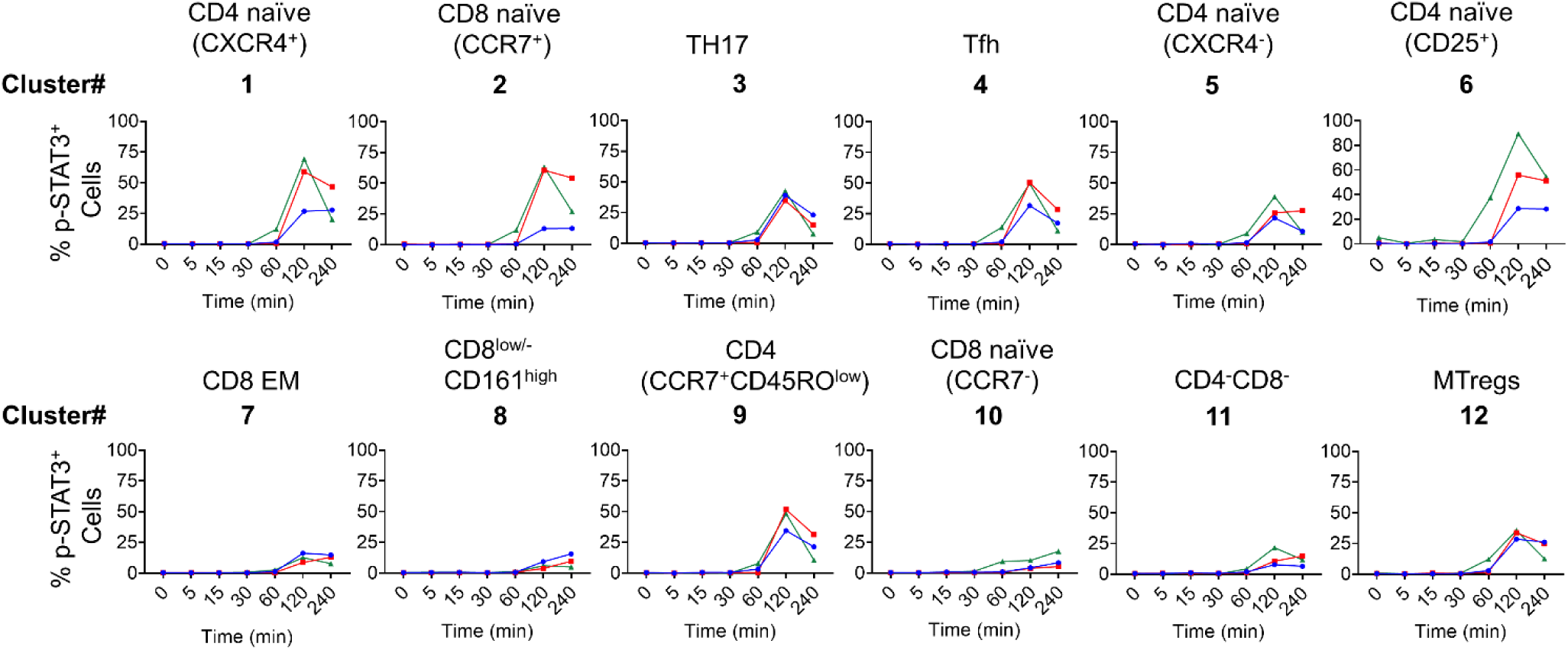
Prolonged IL-1β stimulation induces expression of p-STAT3. **a**, Time-dependent changes in the frequency of p-STAT3^+^ cells within T cell clusters. Red, blue and green lines indicate 3 donors.

### Anakinra abrogates IL-1β-induced signaling activation

Anakinra is a recombinant non-glycosylated IL-1R antagonist, which competitively binds to IL-1R1 and prevents IL-1β binding and downstream signaling. To determine the dependence of IL-1β-induced expression of p-NF-kB and p-p38 on IL-1R1, PBMCs were incubated with Anakinra prior to IL-1β stimulation. Anakinra significantly inhibited IL-1β-induced expression of p-NF-kB in the CCR6^+^ EM T cells and CD11c mDCs and led to trending inhibition in CM and Lin^−^CD161^+^CD25^+^ cells (Fig. 7a-d).

**Fig 7.**
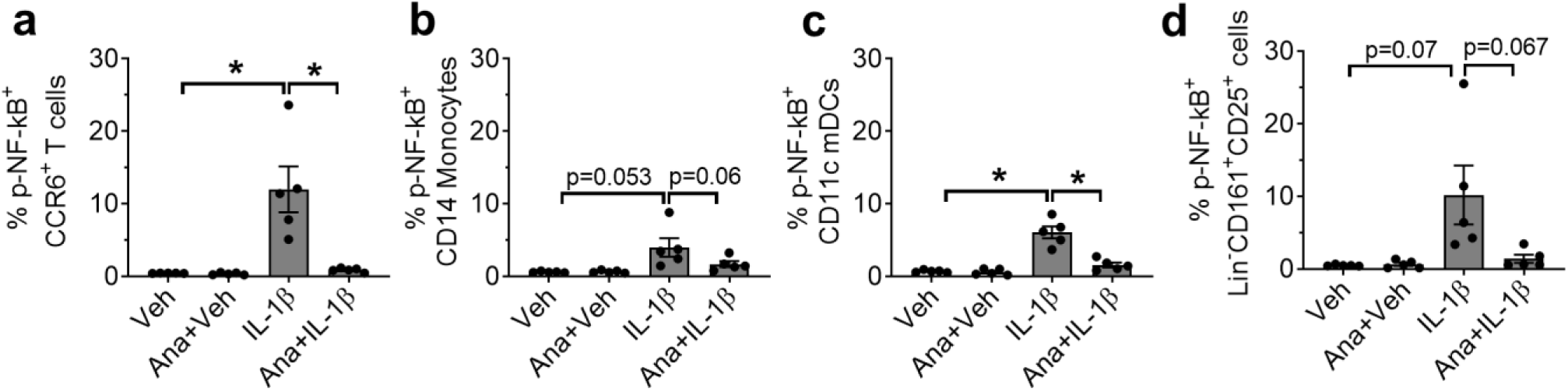
Abrogation of IL-1β-induced signaling by Anakinra (Ana). PBMCs were incubated with 10 µg/ml Anakinra for 30 minutes prior to treatment with vehicle and IL-1β for 15 minutes. Cells were PFA-fixed, washed, barcoded and stained with cell surface and intracellular phospho antibodies. Debarcoded, normalized CyTOF data was analyzed in Flowjo. Frequency of p-NF-kB^+^ **a**, CCR6^+^ T cells, **b**, CD14 monocytes **c**, CD11c mDCs and **d**, Lin^−^CD161^+^CD25^+^ cells. Data shown is mean ± SEM. Unpaired t-test. N=5; * mean ± SEM; p ≤ 0.05.

## Discussion

Inhibition of IL-1β has emerged as a promising therapeutic approach for a variety of inflammatory diseases ^32–35^ and a potential therapy for reducing the cytokine strom associated with COVID-19 ^2,36^. While studies over the past decades have provided significant insights into pathophysiological mechanisms that lead to dysregulated production of IL-1β, knowledge about the specific human immune cell types that IL-1β targets is still limited. We have identified the key human immune cell subtypes that are most vulnerable to IL-1β, as well as the subset-specific signaling pathways that are activated.

CD4 EM T cells were the circulating immune cell subtype most robustly activated by IL-1β. CD4 EM T cells display rapid effector functions as compared to their naïve counterpart and are implicated in various inflammatory diseases including atherosclerosis, rheumatoid arthritis (RA) and diabetes. Increased frequency of CD4 EM T cells are associated with increased disease activity and severity ^37–44^. Interestingly, a recent study demonstrated that T cell intrinsic IL-1R signaling acts as a licensing signal for cytokine production by memory CD4 T cells and that this critical cue is conserved across all EM lineages ^17^. IL-1β enhanced post-transcriptional stability of effector cytokines such as TH17 cytokines, IL-5 and IL-13 via the p38 MAPK pathway in the murine CD4 EM T cells ^17^. Other studies, mostly murine, also demonstrated that IL-1β is crucial for proinflammatory cytokine production by CD4 EM T cells through activation of NF-kB, p38 and JNK ^15,16,18–21^. Consistent with these studies, our CyTOF analyses in human immune cells demonstrated IL-1β-induced signaling predominantly in the CD4 EM subsets including TH17, Tfh and MTregs. IL-1β-induced expression of p-NF-kB was more robust as compared to p-p38 expression. Our data suggests that NF-kB inflammatory pathway may play a more prominent role in IL-1β-mediated regulation of functions of the CD4 EM T subsets in humans. Murine CD4^+^CXCR5^hi^CD25^−^ Tfh cells express IL-1R1, and IL-1β triggers in vivo expansion of these Tfh cells and induces production of IL-4 and IL-21, the two major B cell activating cytokines ^45^. To our knowledge, our study is the first to demonstrate that IL-1β induces NF-kB activation in the human Tfh cells, potentially providing new mechanistic insight into IL-1β-mediated dysregulated humoral response in patients with autoimmune diseases^46^.

Interestingly, IL-1β induced p-NF-kB expression in the EM T cell cluster 8 comprised of two distinct cell subsets, CD4^−^CD8^+^ and CD4^−^CD8^−^ (**Supplementary Fig. 7**) that express high levels of CD161, a marker expressed predominantly on NK cells. CD4^−^CD8^+^CD161^high^ T cells with a similar phenotype as the CD4^−^CD8^+^ cells within cluster 8 are present in the human peripheral blood and represent mucosal-associated invariant T (MAIT) cells ^47–51^. These CD4^−^CD8^+^CD161^high^ T cells express reduced levels of CD8 and share phenotypic features with the Th17 cells, including expression of the chemokine receptors (CCR6, CCR2 and CXCR6), cytokine receptors (IL-23R and IL-18R) and RORC transcription factor ^47,48,51^. Moreover, the CD4^−^CD8^+^CD161^high^ cells belong to the conventional TCR-αβ EM (CD45RO^+^CCR7^−^) T cells ^47,48^ that are not exhausted or terminally differentiated, and a subset of these cells express IL-17, IFN-γ, TNF-α and IL-22 ^47,52^. Notably, gating total CD8 T cells following the strategy used by Billerbeck et al. 47, we were able to distinctly identify the CD161^high^ cells that expressed relatively lower levels of CD8 (**Supplementary Fig. 7e**). Furthermore, the CD4^−^CD8^+^CD161^high^ cells in cluster 8 colocalized with the CD8^low^CD161^high^ subset when overlaid on the total CD8 T cells (**Supplementary Fig. 7f**). In the present study, we did not measure expression of CXCR6, IL-23R and IL-18R. However, the expression of markers including CCR2, CCR6, CD45RO and CD127 on the CD8^+^CD161^high^ T cells in cluster 8 (Fig. 2c) and absence of CCR7 expression (**Supplementary Fig. 7g-m**) suggests that they may be CD8^+^CD161^high^ MAIT cells as reported earlier ^47,48,51^. Further supporting this notion is the study by Turtle et al., which demonstrated that IL-1β in cooperation with T cell receptor signaling enhances cell proliferation and IL-17 secretion in human CD8α^+^CD161^high^ cells ^52^. While MAIT cells play an important role in protective immunity to pathogens, emerging evidence support their pathogenic role in chronic inflammatory, metabolic and autoimmune pathologies ^53–59^. MAIT cells express high levels of tissue-targeted chemokine receptors ^50^ and display an activated proinflammatory TH17 phenotype ^59^. Indeed, increased frequencies of MAIT cells have been reported at the site of inflammation in patients with RA, multiple sclerosis, diabetes, obesity, inflammatory bowel disease and others ^56,57,60,61^.

In addition to the CD4^−^CD8^+^CD161^high^ cells, the CD4^−^CD8^−^ subset within cluster 8 also showed induced expression of p-NF-kB in response to IL-1β (**Supplementary Fig. 7a-c**). These CD4^−^CD8^−^ cells expressed comparable levels of CD161 and displayed a phenotype very similar to the CD4^−^CD8^+^CD161^high^ cells within cluster 8 (**Supplementary Fig. 7a,d and i-m**). Notably, Moreira-Teixeira et al. ^62^ demonstrated that co-stimulation with IL-β, TGF-β and IL-23 induces IL-17 production in the CD4^−^CD161^+^ human invariant NKT (iNKT) cells ^62^. Different cell types including, conventional Th17 cells as well as the unconventional subsets, γδ, NKT and MAIT cells ^50,63–65^ express CD161, and confer pathogenic roles in inflammatory diseases ^53,66,67^. Further work is warranted to establish the identity of the CD4^−^CD8^−^CD161^high^ and CD4^−^CD8^+^CD161^high^ cells (cluster 8) that showed IL-1β-induced expression of p-NF-kB and p-p38, and investigate how IL-1β modulates their effector functions.

Another novel finding of the present study is the identification of CCR6 as a marker to identify T cells most responsive to IL-1β stimulation. Our analysis demonstrated that the IL-1β responsive CCR6^+^ cells belong to the EM phenotype and a majority of these CCR6^+^ EM T cells co-express CCR2 and CD26. Notably, circulating human CD4 and CD8 T cells capable of producing IL-17 are enriched in the CCR6^+^ memory T cells ^68^, and CD4 helper T cells that produce type 17 cytokines express the highest levels of the catalytically active dipeptidylpeptidase IV (CD26) ectoenzyme ^29^. Interestingly, while Tregs are largely anti-inflammatory, a proportion of the circulating MTregs in healthy humans express high levels of RORC and secrete IL-17 ^69^. IL-1β together with IL-23 plays a critical role in human TH17 cell development and regulation of TH17 cytokines ^70–73^. It is possible that the cells within the CCR6^+^ EM T cell population that respond to IL-1β bear a TH17-like characteristic. Abundant evidence implicate TH17 cells and sustained production of IL-17 in the pathogenesis of diverse inflammatory and autoimmune diseases ^74–80^. Increased frequency of TH17 cells and higher levels of TH17 cytokines have been reported in patients with inflammatory diseases and blocking the IL-17 pathway has shown potential benefit in some inflammatory diseases ^74,76,78,81,82^. IL-17 confers pathogenic effects through enhanced production of cytokines (IL-6, IL-1β, TNF-α, GM-CSF), chemokines (IL-8, CXCL1, CXCL2, CCL20, MCP-1), metalloproteinases and other inflammatory molecules ^83^. These proinflammatory mediators promote inflammation, immune cell infiltration and tissue damage. Additionally, IL-17-induced IL-6 maintains the TH17-T cell population ^83^. The IL-1β-responsive EM T cells expressing CCR6, CCR2 and possibly other chemokine receptors may be recruited to sites of inflammation where IL-1β together with other proinflammatory cytokines such as IL-23 or antigen-mediated TCR activation may activate them to produce proinflammatory cytokines and cause tissue damage. Taken together, findings raise the interesting possibility that CCR6^+^ EM T cells and/or IL-17 producing cells likely plays an important role in mediating IL-1β-induced inflammatory response in humans. One probable anti-inflammtory mechanism of IL-1 blockers may thus be via reduction of the proinflammatory TH17 response. Indeed, in RA patients, Anakinra therapy associated with clinical improvement decreased the frequency of TH17 cells and serum levels of IL-17 and IL-21^84^.

Monocytes and DCs represent major producers of IL-1β and autocrine signaling by IL-1β is known to induce its own production in human monocytes ^85^. All three conventional human monocyte subsets namely, CM, IMM and NCM produce IL-1β. However, NCM produced lower levels of IL-1β, both under basal conditions and after LPS stimulation as compared to the CM and IMM ^86^. It is unclear if all monocyte subsets respond to IL-1β stimulation. Our CyTOF analysis identified 8 monocyte subsets and CD11c mDCs and CD123 pDCs. Interestingly, our results demonstrate that IL-1β induces expression of p-NF-kB and p-p38 only in the CM subsets. Human monocyte subsets display remarkable heterogeneity in their surface marker expression, effector functions and differentiation potential ^87^. CM exhibit a more proinflammatory phenotype as compared the NCM and IMM subsets. Our data showing different responsiveness to IL-1β stimulation further adds to the knowledge of the unique characteristics of the monocyte subsets. Within the two DCs subsets, IL-1β-induced p-NF-kB expression was seen only in the CD11c mDCs. Notably, unlike T cells and NK cells, IL-1β-induced expression of p-NF-kB, but not of p-p38 in the CM and CD11c mDCs remained elevated for prolonged time periods. This data suggests involvement of the NF-kB pathway in the autocrine IL-1β signaling that may lead to continued production of IL-1β under chronic inflammatory conditions.

Within the NK subsets, IL-1β induced expression of p-NF-kB in the CD16^−^CD56^dim^CD161^+^ (cluster 2) and CD16^−^CD56^bright^CD161^−^ cells (cluster 3). CD161 is a C-type lectin receptor and CD161 expression divides circulating NK cells in healthy adults into two distinct populations, with ~80% cells being CD161^+ 88^. Both CD56^bright^ and CD56^dim^ subsets comprise CD161^+^ cells. Notably, CD161^+^ NK cells within the CD56^dim^ subset produced more IFN-γ than their CD161^−^ counterpart, were more proliferative and showed an increased responsiveness to IL-12 and IL-18 ^88^. Furthermore, a higher frequency of the CD161^+^ NK cells were present in the inflamed intestinal lamina propria of patients with inflammatory bowel disease ^88^. Here, using the high-dimensional approach, we have identified a subset of CD161^+^ NK cells with dim expression of CD56 that responds to IL-1β stimulation. It would be interesting to investigate if our cluster 2 shares phenotypic and functional properties with that of the CD161^+^ NK cells identified by Kurioka et al ^88^ or constitute a separate subset that responds specifically to IL-1β.

Intriguingly, the only cell type where the majority of the cells showed expression of p-NF-kB after IL-1β stimulation was a subset of CD161^+^ cells (cluster 17) that were Lin^−^(CD3^−^CD4^−^CD8^−^CD19^−^CD14^−^CD16^−^HLA^−^DR^−^CD123^−^) and expressed much higher levels of CD25 than the CD56^bright^ NK cells (cluster 3) (**Supplementary Figs. 10b,c**). Moreover, the majority (~80%) of the cells in cluster 17 were CD56^−^. Interestingly, cluster 17 showed a robust induction of p-NF-kB, p-p38 as well as activation of STAT1, STAT3 and STAT5 at later time points. Human innate lymphoid cells (ILCs) also express CD161 ^89^ and Ohne et al. ^22^ demonstrated that IL-1β plays a critical role in inducing proliferation, cytokine production, maturation and plasticity of human ILC2. IL-1β mediated these effects in combination with IL-2 and/or IL-12, and NF-kB inhibition blocked IL-1β-induced responses ^22^. Notably, p-NF-kB^+^ cells within our cluster 17 bear resemblance to the phenotype of the ILC2. Consistent with the higher expression of CD127 in the ILC2 cells as compared to the CD56^bright^ NK cells as shown by Ohne et al., ^22^ the p-NF-kB^+^ cells within cluster 17 expressed higher CD127 than the p-NF-kB^+^ cells within the CD56^bright^ NK cells (cluster 3) (**Supplementary Fig. 10d)**. However, in contrast to the ILC2, around 20-30% of the p-NF-kB^+^ cells within cluster 17 were found to be CD56^dim^ (**Supplementary Figs. 10e,f**).

IL-1β is an apical proinflammatory cytokine that exacerbates inflammation through activating both innate and adaptive immune cells and production of a variety of proinflammatory cytokines and chemokines. Indeed, induction of p-STAT1, p-STAT3 and p-STAT5 expression across multiple immune cell subtypes after prolonged IL-1β stimulation indicates signaling mediated by secondary mediators such as IL-6, IFN-γ, IL-17A. Robust induction of p-NF-kB expression in the CCR6^+^ EM T cells, CD11c mDC and the Lin^−^CD161^+^CD25^+^ cells indicate that these cell types may play an important upstream role in IL-1β-mediated inflammation including cytokine storm.

This novel comprehensive atlas of IL-1β-induced signaling in circulating human immune cells provides new insights into the cell types most vulnerable to IL-1β activation and the timing of these effects in humans. Together, these provide key data for the study and design of more targeted approaches to chronic inflammatory diseases that may lessen adverse effects. Moreover, our work has clear implications for the development of immune phenotypic approaches to identify COVID-19 subjects at higher risk of cytokine storm and those most likely to benefit from therapies targeting IL-1β or its downstream mediators such as IL-6.

## Acknowledgements

We thank Mike Solga and Claude Chew from the UVA Flow Cytometry Core for their excellent technical assistance.

## Sources of Funding

This work was supported by the American Heart Association Innovative Project Award and R01HL 136098 (CAM).

## Disclosures

None

## Non-standard abbreviations and acronyms

CyTOF: Mass Cytometry
IL-1β: Interleukin-1 beta
EM: Effector memory
Mtregs: Memory T regulatory cells
CM: Classical monocytes
NCM: Non-classical monocytes
IMM: Intermediate monocytes
mDC: Myeloid dendritic cells
pDC: Plasmacytoid dendritic cells
NK: Natural Killer
IL-1R: Interleukin-1 receptor
PBMC: Peripheral blood mononuclear cells
PFA: Paraformaldehyde
CSB: Cell staining buffer
SPADE: Spanning-tree Progression Analysis of Density-normalized Events
UMAP: Uniform Manifold Approximation and Projection

## Notes

### Competing Interest Statement

The authors have declared no competing interest.

