## Supplementary table and figures for "Identification of Human Immune Cell Subtypes Most Vulnerable to IL-1β-induced Inflammatory Signaling Using Mass Cytometry"

**Supplementary Table 1.** CyTOF antibodies.

| Metal Tag | Clone | Manufacturer | Antigen |
|---|---|---|---|
| **Surface Markers** | | | |
| 89Y | H130 | Fluidigm | CD45 |
| 141 Pr | G034E3 | Fluidigm | CCR6 |
| 142 Nd | HIB19 | Fluidigm | CD19 |
| 143 Nd | 6H6 | Fluidigm | CD123 |
| 144 Nd | H-B7 | Biolegend | CD38 |
| 145 Nd | RPA-T4 | Fluidigm | CD4 |
| 146 Nd | ICRF44 | Biolegend | CD11b |
| 147 Sm | 5-271 | Biolegend | CD36 |
| 148 Nd | RM052 | Fluidigm | CD14 |
| 149 Sm | 2A3 | Fluidigm | CD25 |
| 151 Eu | 34141 | R & D Systems | IL-1R2 |
| 152 Sm | 732229 | R & D Systems | IL-1R1 |
| 154 Sm | MU5UBEE | Affymetrix | CXCR5 |
| 159 Tb | HP-3G10 | Fluidigm | CD161 |
| 161 Dy | BA5b | Fluidigm | CD26 |
| 162 Dy | Bu15 | Fluidigm | CD11c |
| 163 Dy | G025H7 | Fluidigm | CXCR3 |
| 164 Dy | UCHL1 | Fluidigm | CD45RO |
| 165 Ho | A019D5 | Fluidigm | CD127 |
| 167 Er | G043H7 | Fluidigm | CCR7 |
| 168 Er | SK1 | Fluidigm | CD8 |
| 170 Er | UCHT1 | Fluidigm | CD3 |
| 172 Yb | MHM-88 | Fluidigm | IgM |
| 173 Yb | K036C2 | Biolegend | CCR2 |
| 174 Yb | L243 | Fluidigm | HLA-DR |
| 175 Lu | 12G5 | Fluidigm | CXCR4 |
| 176 Yb | N901 | Fluidigm | CD56 |
| 209Bi | 3G8 | Fluidigm | CD16 |
| **Intracellular Signaling Proteins** | | | |
| 150 Nd | 47 | Fluidigm | pSTAT5[Y694] |
| 153 Eu | 58D6 | Fluidigm | pSTAT1 [Y701] |
| 155 Gd | SB40a | Novus Biologicals | Tollip |
| 156 Gd | D3F9 | Fluidigm | P-p38[T180/Y182] |
| 158 Gd | 4/P-Stat3 | Fluidigm | pSTAT3 [Y705] |
| 166 Er | K10-895.12.50 | Fluidigm | P-NFκBp65[S529] |
| 171 Yb | D13.14.4E | Fluidigm | P-ERK1/2[T202/Y204] |

**Supplementary Fig 1. PFA fixation affects cell surface marker epitopes and antibody reactivity.** Cryopreserved PBMCs were either fixed or not in 1.6% PFA before staining with the CyTOF antibody panel. Samples were run on CyTOF and data analyzed by Flowjo. Representative data from a donor is shown.

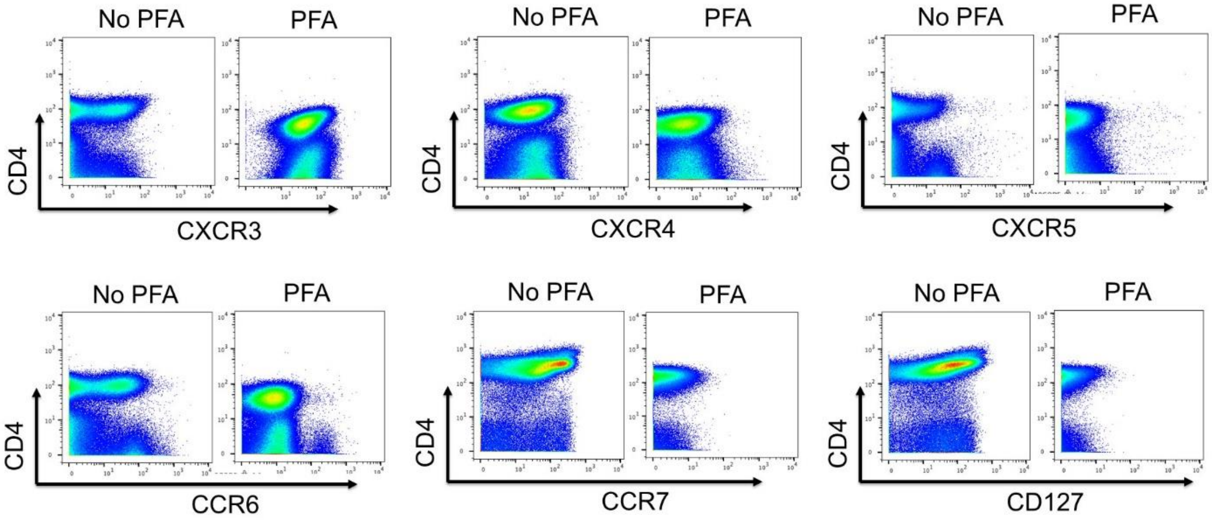


**Supplementary Fig 2. Antibody staining simultaneously during IL-1β stimulation does not interfere with IL-1β-induced signaling.** PBMCs were rested for one hour in culture media at 37°C. A small volume of a cocktail of surface antibodies (CD127, CXCR3, CXCR4, CXCR5, CCR6 and CCR7) was added to PBMCs during IL-1β stimulation for 15 minutes at 37°C. Cells were immediately fixed with PFA, washed and stained with the remaining cell surface antibodies followed by labeling with the intracellular phospho markers. Flowjo plots from a representative donor is shown.

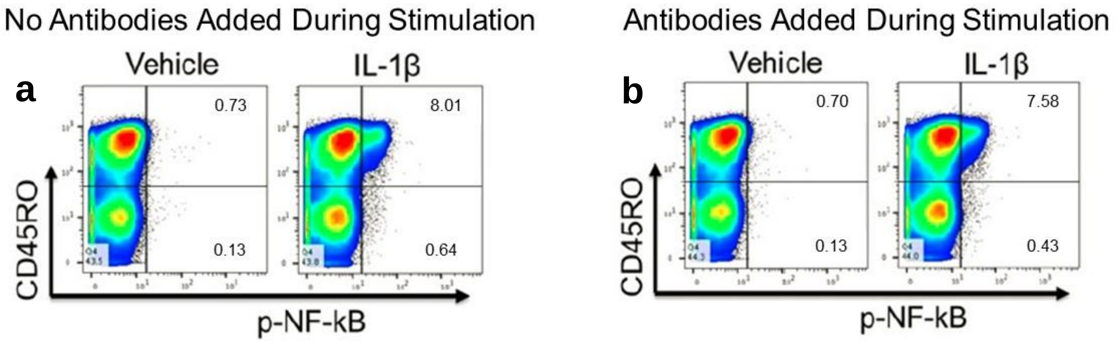

**Supplementary Fig 3. Gating Scheme.** Debarcoded normalized .fcs files were analyzed in Flowjo. **a,** Iridium staining ($^{191}$Ir DNA intercalator) identified cells, and intact singlets were discerned from debris and cell aggregates by gating on cells with higher DNA content. **b,** Sample-specific stringency adjustment by manual biaxial plotting on normalized barcode separation distance and mahalonobis distance was performed to ensure accurate sample identity. **c,** CD45$^+$ cells were separated into CD3$^+$ T cells, CD19$^+$ B cells and CD3$^-$CD19$^-$ cells. **d,** Contaminating monocytes were further removed from the CD3$^+$ T cells before exporting .fcs files for T cell clustering. **e,** CD3$^-$CD19$^-$ cells were divided into HLA-DR$^-$ and HLA-DR$^+$ cell populations to be used for clustering NK cells and monocytes/DCs, respectively. **f-j,** Representative dot plots showing gating strategy for quantitating frequency of phospho$^+$ cells within clusters. A gate in the vehicle-treated samples for individual donor with a threshold of 0.5-1% was made and was then applied to the IL-1β-stimulated samples.

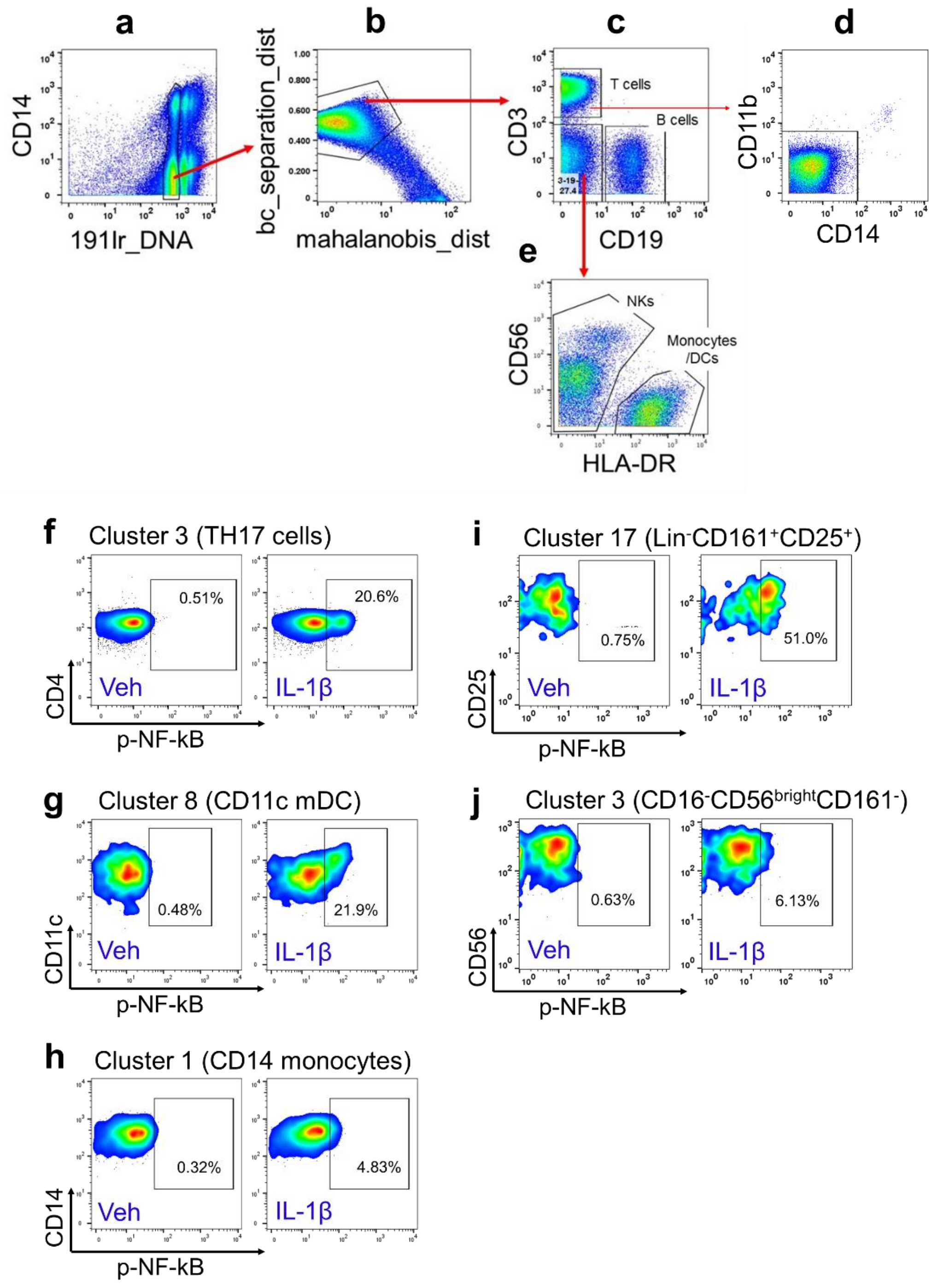

**Supplementary Fig 4. IL-1β Dosimetry.** PBMCs were stimulated with varying concentrations of IL-1β for 15 minutes, fixed with PFA and stained with cell surface and intracellular phospho markers. Frequency of **a and b,** p-NF-kB⁺ and **c and d,** p-p38⁺ cells within CD4 memory (CD45RO⁺) cells and CD11c mDCs.

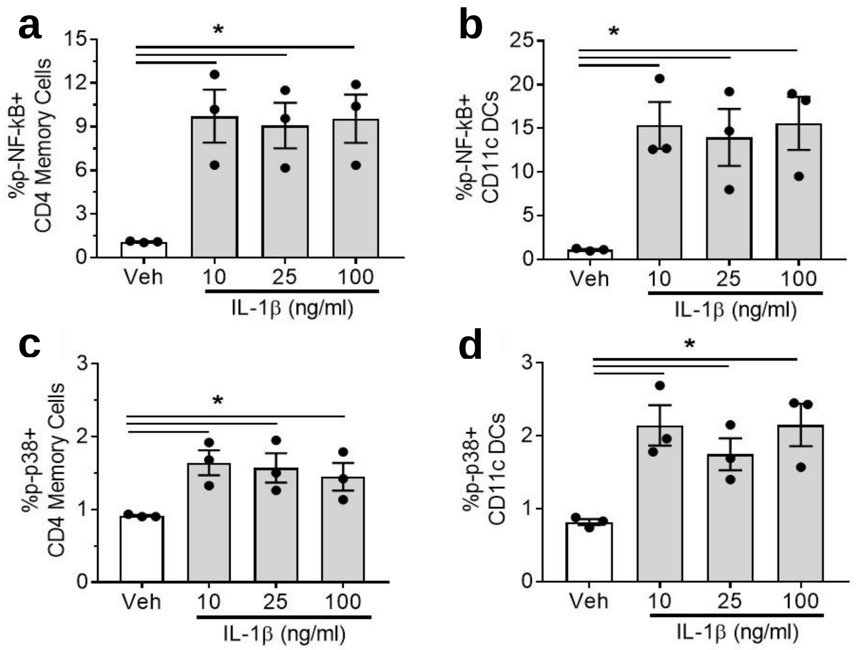

**Supplementary Fig 5. IL-1β-induces prolonged expression of p-NF-kB in classical monocytes (CM) and CD11c myeloid dendritic cells (mDC) but transient induction in memory T cells, NK cells and Lin$^-$CD161$^+$CD25$^+$ cells**. Time-dependent changes in the frequency of **a,** p-NF-kB$^+$ and **b,** p-p38$^+$ cells within clusters of T cells, monocytes, DCs, NK cells and Lin$^-$CD161$^+$CD25$^+$ cells. Red, blue and green lines represent 3 donors. ** Note: scale for the Lin$^-$CD161$^+$CD25$^+$ is different.

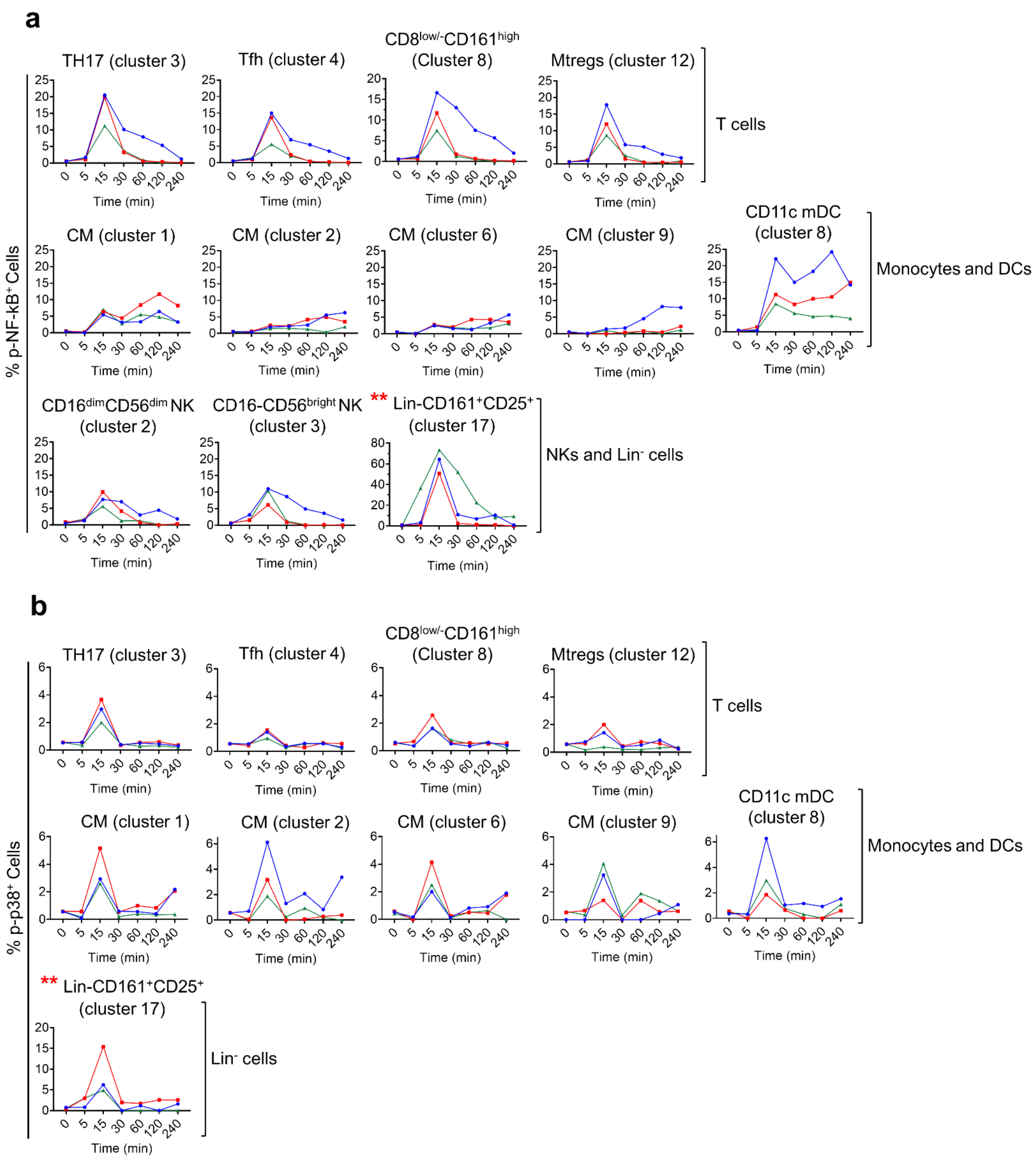

**Supplementary Fig 6. Time-dependent changes in the frequency of p-STAT1⁺, p-STAT3⁺ and p-STAT5⁺ cells within different clusters of T cells, monocytes, DCs and within the Lin-CD161⁻CD25⁺ cells.** Red, blue and green lines indicate 3 donors.

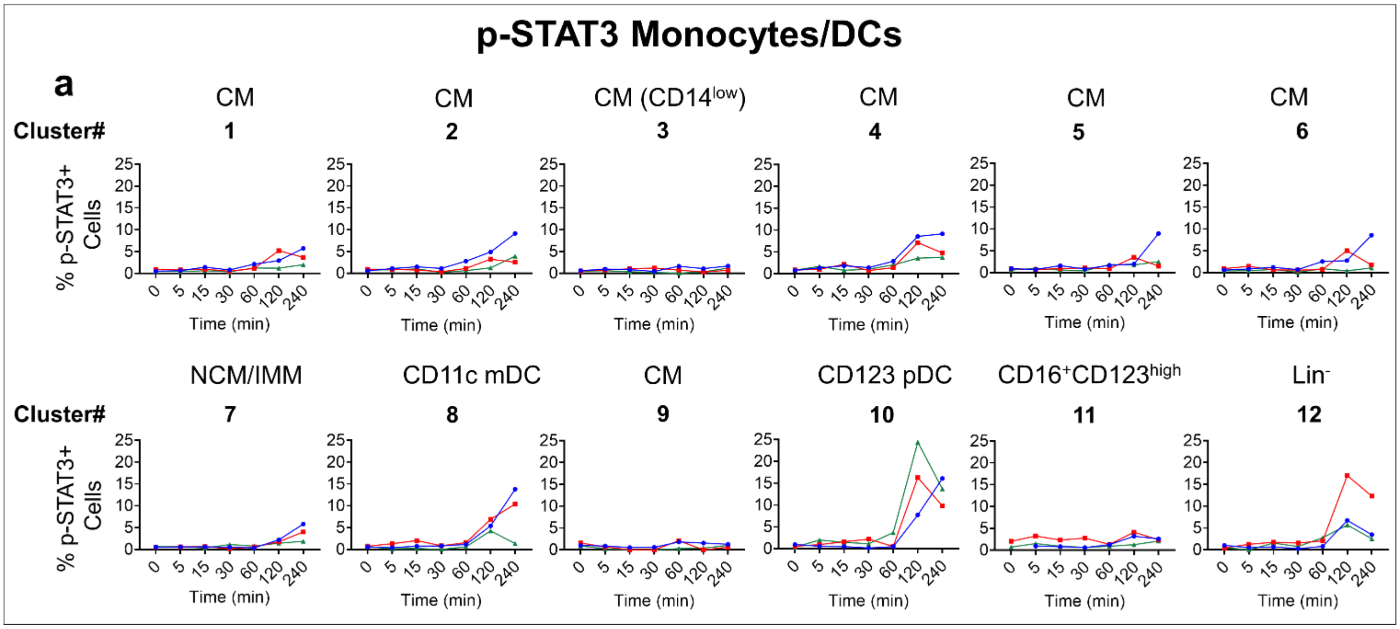


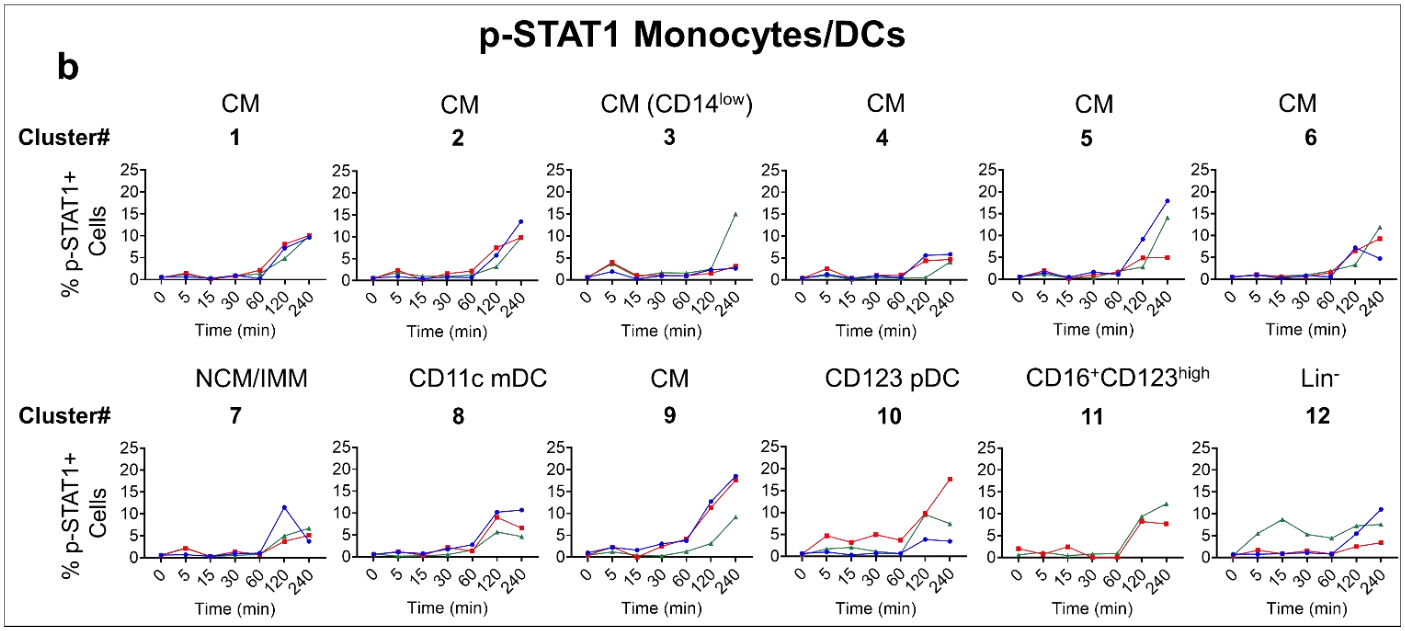

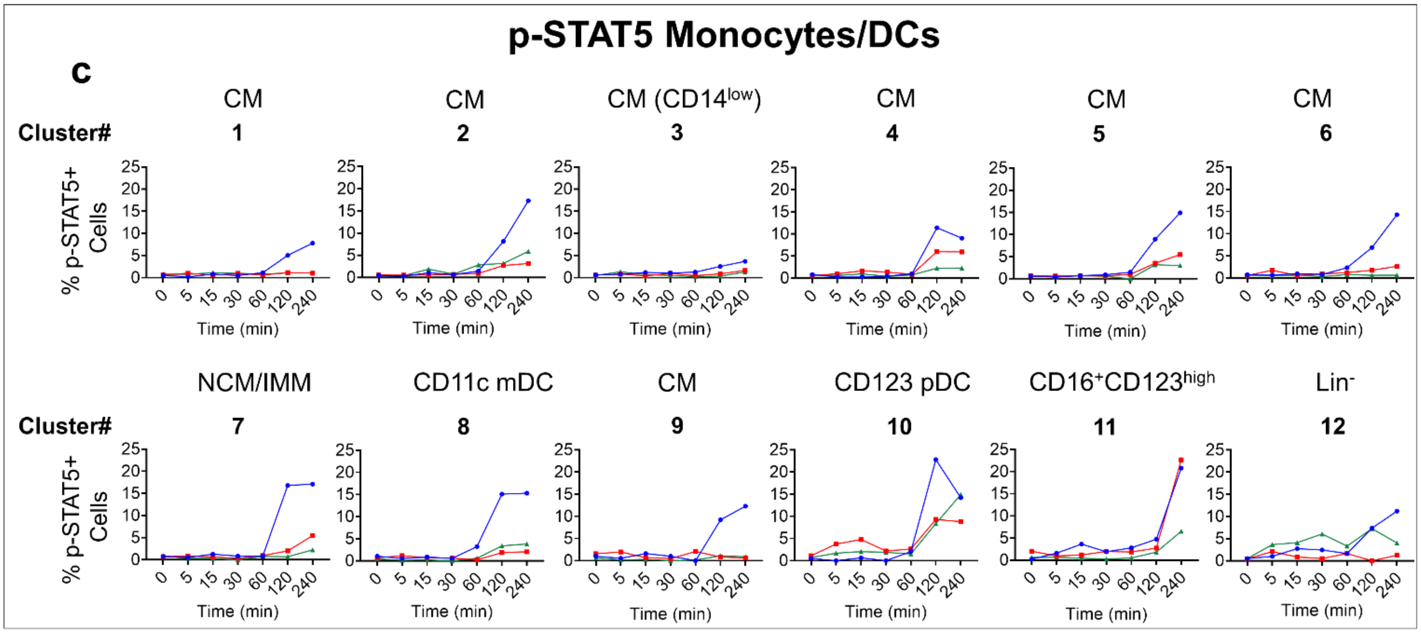
p-STAT5 Monocytes/DCs
c
Cluster#
CM 1
CM 2
CM (CD14low) 3
CM 4
CM 5
CM 6
NCM/IMM 7
CD11c mDC 8
CM 9
CD123 pDC 10
CD16+CD123high 11
Lin- 12
% p-STAT5+ Cells
Time (min)
0 5 15 30 60 120 240
0 5 10 15 20 25

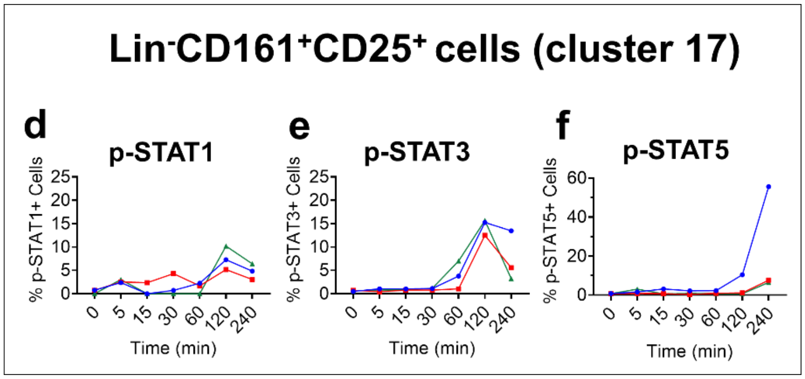
Lin-CD161+CD25+ cells (cluster 17)
d
p-STAT1
% p-STAT1+ Cells
e
p-STAT3
% p-STAT3+ Cells
f
p-STAT5
% p-STAT5+ Cells
Time (min)
0 5 15 30 60 120 240

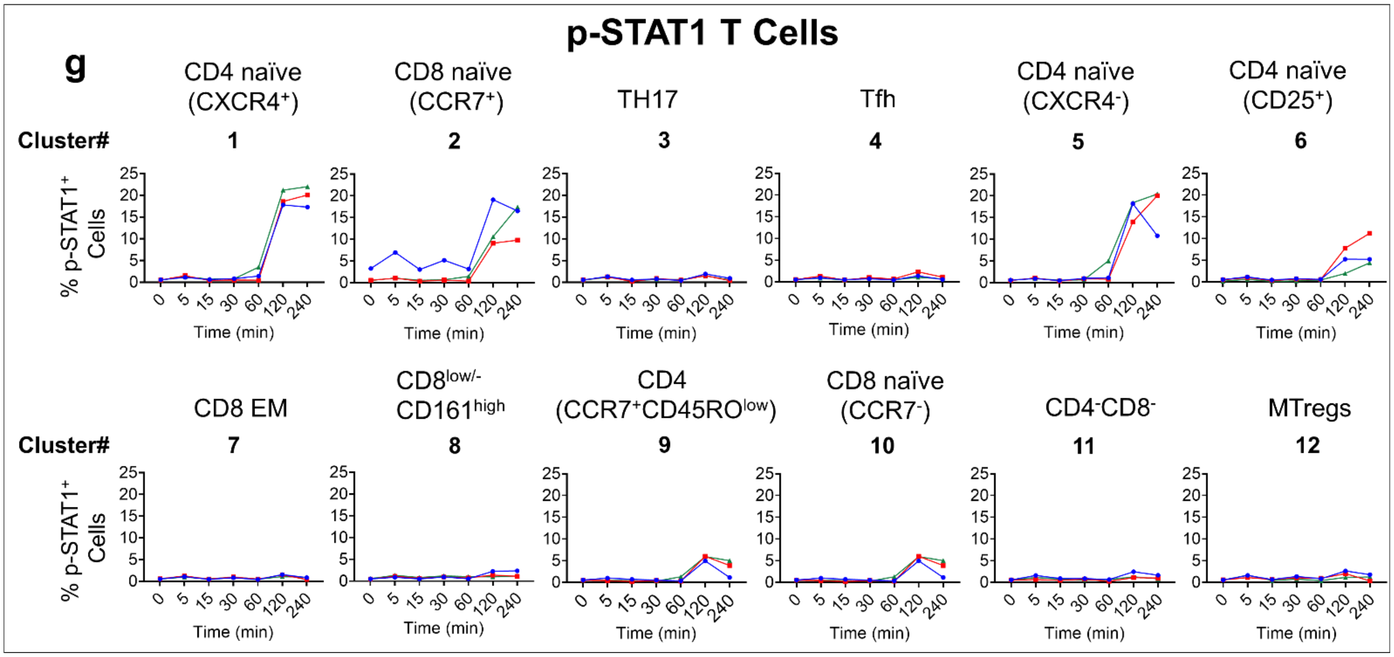
p-STAT1 T Cells
g
Cluster#
CD4 naïve (CXCR4+) 1
CD8 naïve (CCR7+) 2
TH17 3
Tfh 4
CD4 naïve (CXCR4-) 5
CD4 naïve (CD25+) 6
CD8 EM 7
CD8low/- CD161high 8
CD4 (CCR7+CD45ROlow) 9
CD8 naïve (CCR7-) 10
CD4-CD8- 11
MTregs 12
% p-STAT1+ Cells
Time (min)
0 5 15 30 60 120 240

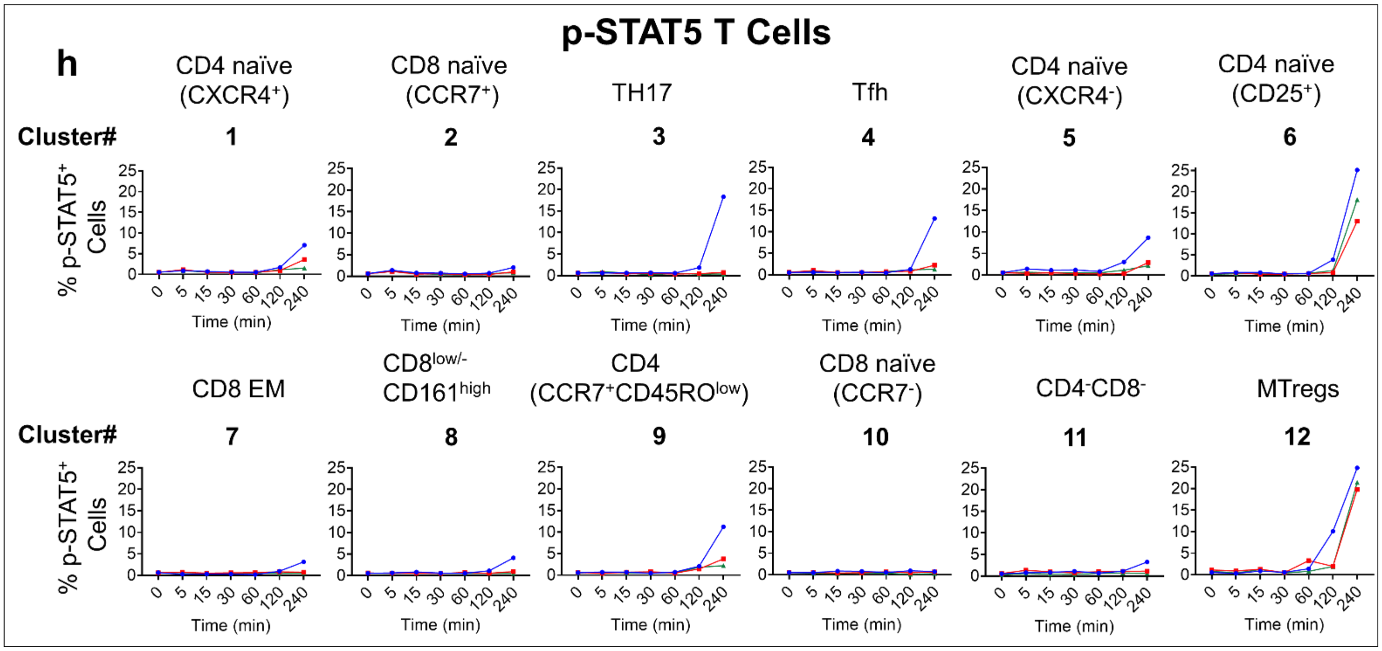
p-STAT5 T Cells
h
CD4 naïve (CXCR4⁺)
CD8 naïve (CCR7⁺)
TH17
Tfh
CD4 naïve (CXCR4⁻)
CD4 naïve (CD25⁺)
Cluster#
1
2
3
4
5
6
% p-STAT5⁺ Cells
Time (min)
CD8 EM
CD8low/- CD161high
CD4 (CCR7⁺CD45ROlow)
CD8 naïve (CCR7⁻)
CD4⁻CD8⁻
MTregs
Cluster#
7
8
9
10
11
12
0 5 15 30 60 120 240
0 5 10 15 20 25

**Supplementary Fig 7. T cell cluster 8 contains two distinct populations (CD8$^-$ and CD8$^+$) that both express high levels of CD161. a,** Representative biaxial gating of cluster 8 from one donor showing CD8$^-$ and CD8$^+$ cells with comparable expression of CD161. **b,** Frequency of CD8$^-$ and CD8$^+$ cells within cluster 8. **c,** Frequency of p-NF-kB$^+$ cells within CD8$^-$ and CD8$^+$ subsets. **d,** Histogram overlay depicting CD26 expression in cluster 3 and CD8$^-$ and CD8$^+$ subsets within cluster 8. **e,** Representative 2-D plot of total CD8 T cells from one donor showing distribution pattern of the CD8$^+$ subsets with low and high CD161 expression, and reduced expression of CD8 on the CD161$^{high}$ cells. **f,** Overlay of the cluster 8 CD8$^+$ subset on total CD8 T cells showing colocalization of the former specifically within the CD8$^{low}$CD161$^{high}$ subset. **g and h,** Expression of CCR2 and CCR6 on CD8$^{low}$CD161$^{high}$ cells. **i-m,** Histogram overlays of cluster 8 CD8$^-$ and CD8$^+$ subsets showing similar marker expression.

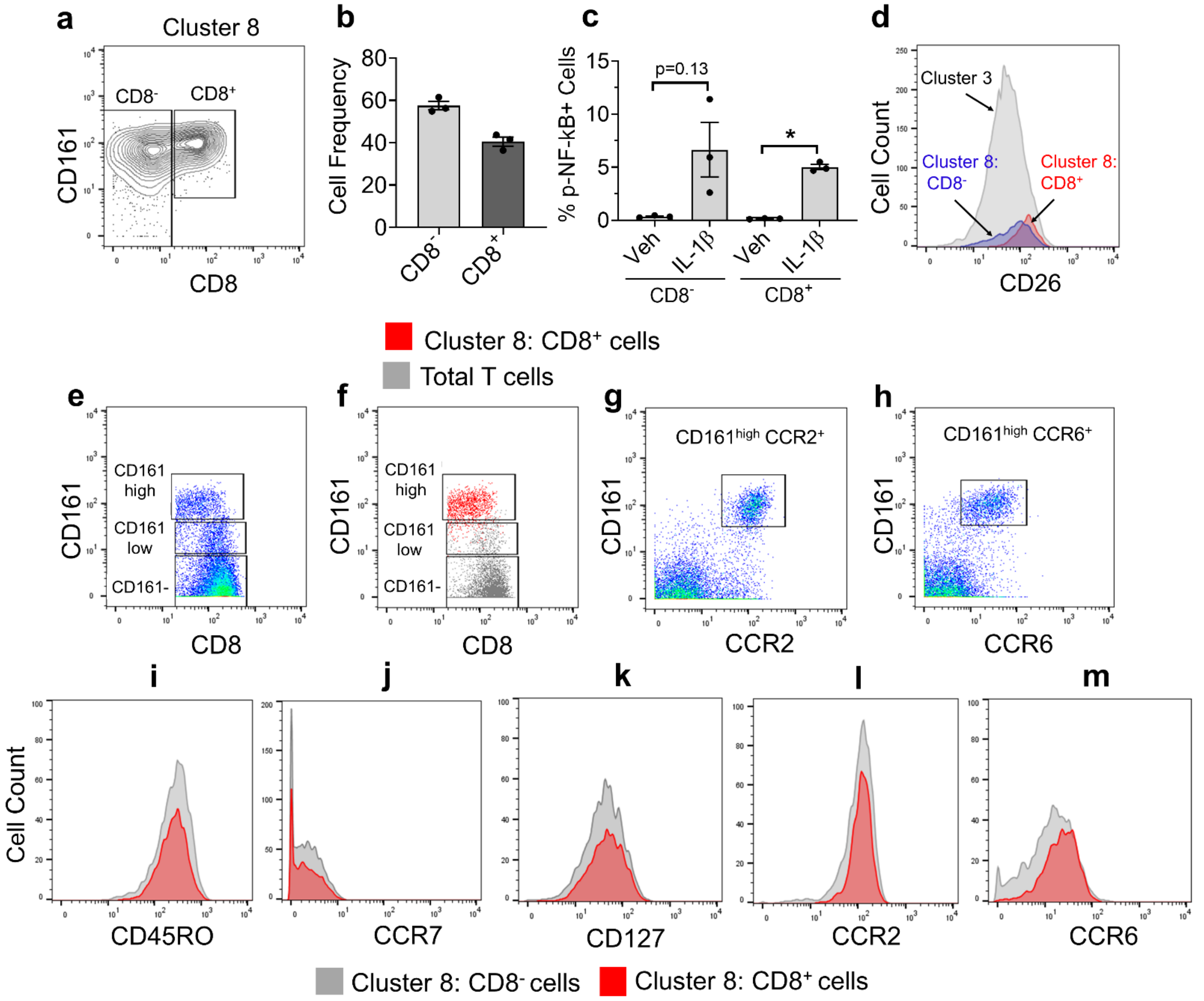

**Supplementary Fig 8. Stimulation with anti-CD3, anti-CD28 and IL-2 increases expression of IL-1R2 on memory CD4 T cells.** Freshly purified PBMCs from 2 healthy donors were stained with anti-IL-1R2 and analyzed by flow cytometry. Alternatively, cells were cultured for 48 hours in the presence of anti-CD3, anti-CD28 and IL-2, and then stained with anti-IL-1R2. Cell surface expression of IL-1R2 on CD4 T cells as analyzed by flow cytometry is shown.

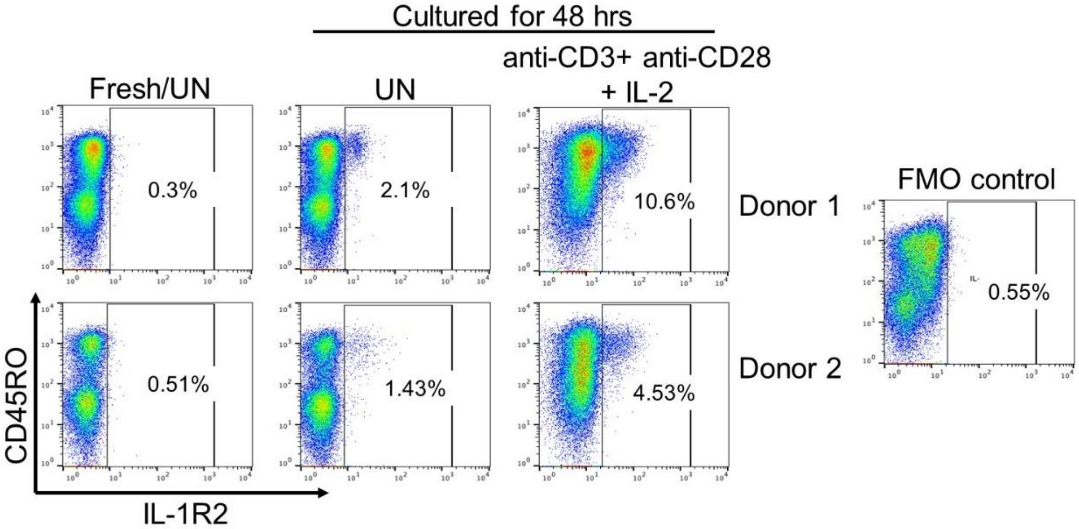

**Supplementary Fig 9. Monocyte cluster 7 was heterogeneous and contained both intermediate and non-classical monocytes.** Dot plots showing that while other clusters such as cluster 1 (CD14 monocytes) and cluster 8 (CD11c mDC) were homogenous, cluster 7 contained intermediate and non-classical monocytes and also $CD14^{-}CD16^{-}$ cells.

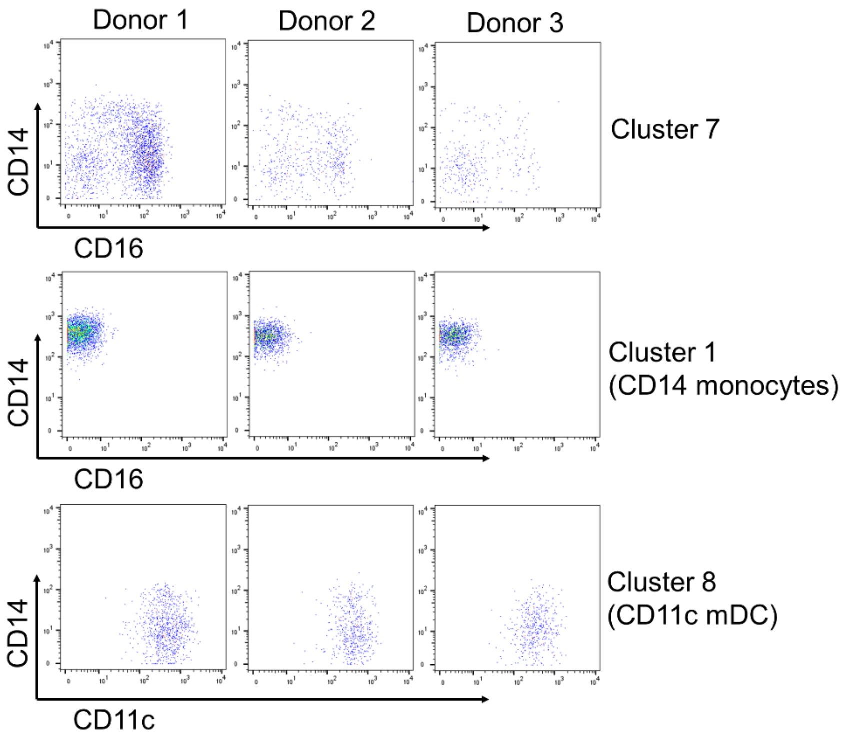

**Supplementary Fig 10. Cluster 17 (Lin$^-$CD161$^+$CD25$^+$) cells do not represent T cells and do not express lineage markers. a,** Overlay of cluster 17 cells on total T cells demonstrating lack of expression of CD3, CD4 and CD8 in the CD161$^+$CD25$^+$ cells but comparable expression of CD25 as the CD4 T cells. **b,** Expression of lineage markers CD19, HLA-DR, CD14, CD16, CD123 on cluster 17 cells. Overlay of cluster 17 from 3 donors. **c,** Overlay of clusters 2, 3 and 17 showing CD25 and CD56 expression. **d,** Overlay of p-NF-kB$^+$ cells in clusters 3 (CD56$^{bright}$ NK cells) and 17 in 3 donors showing higher expression of CD127 on p-NF-kB$^+$ cells in cluster 17 as compared to the p-NF-kB$^+$ cells on the CD56$^{bright}$ NK cells. **e,** Plot showing p-NF-kB$^+$ cells in cluster 17 overlaid on all CD3$^-$CD19$^-$HLA-DR$^-$ cells. **f,** Frequency of CD56$^{bright}$, CD56$^{dim}$ and CD56- cells within total CD3$^-$CD19$^-$HLA-DR$^-$ cells and cluster 17.

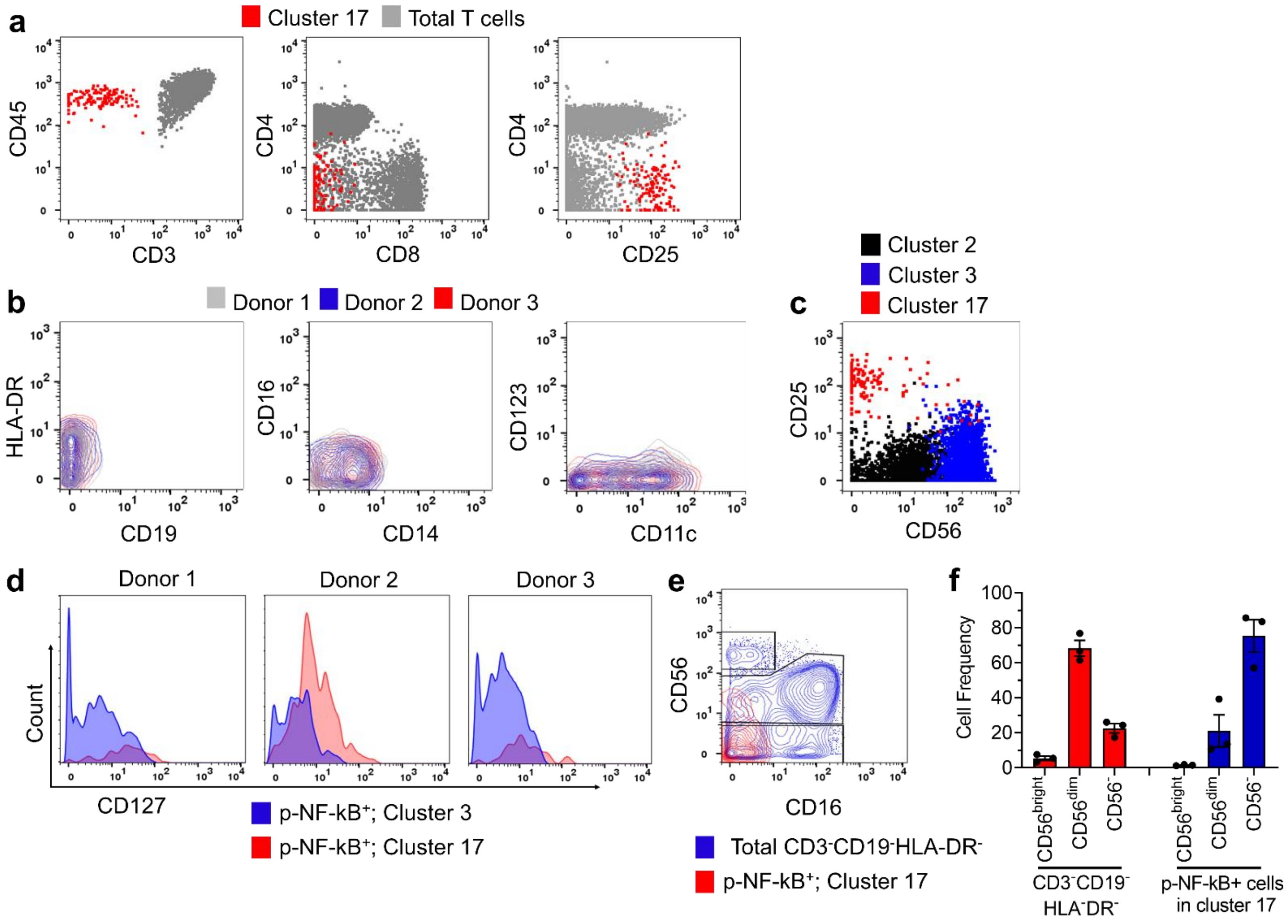
